# Mixed Culture of Bacterial Cell for Large Scale DNA Storage

**DOI:** 10.1101/2020.02.21.960476

**Authors:** Min Hao, Hongyan Qiao, Yanmin Gao, Zhaoguan Wang, Xin Qiao, Xin Chen, Hao Qi

**Author notes:** Equal contribution (Min Hao, Hongyan Qiao, Yanmin Gao). Correspondence should be addressed to H. Q.

## Abstract

DNA emerged as novel material for mass data storage, the serious problem human society is facing. Taking advantage of current synthesis capacity, massive oligo pool demonstrated its high-potential in data storage in test tube. Herein, mixed culture of bacterial cells carrying mass oligo pool that was assembled in a high copy plasmid was presented as a stable material for large scale data storage. Living cells data storage was fabricated by a multiple-steps process, assembly, transformation and mixed culture. The underlying principle was explored by deep bioinformatic analysis. Although homology assembly showed sequence context dependent bias but the massive digital information oligos in mixed culture were constant over multiple successive passaging. In pushing the limitation, over ten thousand distinct oligos, totally 2304 Kbps encoding 445 KB digital data including texts and images, were stored in bacterial cell, the largest archival data storage in living cell reported so far. The mixed culture of living cell data storage opens up a new approach to simply bridge the in vitro and in vivo storage system with combined advantage of both storage capability and economical information propagation.

## 1. Introduction

While being biological material carrying genomic information, DNA has been proven of great potential in storing information in its nucleic acid sequence for long-term in high density. The increased capability of high throughput chip synthesis based writing and next generation sequencing based reading technologies greatly advanced the development of synthesis nucleic acid mediated archival storage. Simply put, information was synthesized into DNA oligo molecule and then read out by sequencing. Till now, a number of systems have been developed storing massive archival data into synthetic oligo pool.^[1, 2]^ Classical electrical communication and computing algorithms such as Fountain and Reed-Solomon code have been adapted for conversation of digital binary information to four letters nucleic acids sequence and error correction.^[3, 4]^ Restricted by current high throughput oligo synthesis techniques, oligo with from 100 to around 200 nts in length was the major materials for information storage in test tube. However, the oligo size well fits with the major commercial sequencing platform, such as Illumina,^[5]^ by which sequence from 50 to 200 nucleotides can be obtained at single read from one oligo terminal end. Furthermore, the cost of chip-based synthesis achieved the lowest DNA synthesis, at least one or two orders of magnitude lower than traditional column-based oligo synthesis. Thus far, as medium materials synthesis oligo pool based in vitro system has been largely expanded for up to 200 MB information storage.^[6]^

Besides test tubes, microbe cells are able to carry the synthesis DNA material with many advanced features for archival information storage. In comparing with the cell-free in vitro system, the genomic maintenance mechanism ensure DNA molecule replicated in a high-fidelity manner in living cells and then higher stability and longer storage period could be expected. Moreover, the DNA molecule copy rate is several orders of magnitude higher than general in vitro replication methods, such as PCR. These advanced features make living cell an attractive materials for copy and distribution of information at low cost. Synthetic DNA fragment encoding archival data have been reported being inserted into genome of various organisms, including *E. coli*,^[7]^ *B. subtilis*^[8]^ and yeast.^[9]^ Molecular tools were developed from engineered various DNA maintenance and genome modification systems, including reverse-transcription,^[10]^ recombinase^[11]^ and CRISPR-cas,^[12]^ for directly writing archival data into genome in a highly controlled fashion. Moreover, circular plasmid was designed for carrying information as well and the multiple copy number of plasmids in microbial cell could facilitate the recovery of DNA material. Seemingly, the in vitro and in vivo DNA storage approach develop as mutually independent system. For in vitro system, massive short piece oligos including even up to 1E10 distinct strands^[13]^ from microchip synthesis were read out by a straight reading workflow comprising PCR amplification and NGS sequencing.^[14]^ In contrast, cell is technically able to store much larger DNA fragment. For a long time, people used to save hundreds of kilobase pairs DNA fragment cloned from human genome in *E. coli* cells.^[15]^ However, being limited by current technological capability, synthesis of large DNA fragment, generally over kilo-nucleotides, is a highly time and cost consuming procedure.^[16]^ Even though entire bacterial chromosome has been synthesized completely,^[17]^ it requires many efforts to carefully design the oligo units and probably takes long time, generally over months, to build them into large fragment.^[18]^ Moreover, it is relatively complicated to efficiently transform large DNA inside cell. Thus far, in vivo DNA storage has only been tested in a relatively small scale, no larger than few thousand nucleotides,^[19]^ far smaller than in vitro system. In considering storage capability, massive short oligo pool has advantage in the ease of scale-up and synthesis cost. However, DNA storage inside cell has advantage in stable DNA material maintaining for long period of time and low cost replication.^[2]^

Here, we demonstrated that mixed culture of bacterial cells carrying massive DNA oligos as economical and sustainable material for stable information storage, in which massive DNA oligos with hundreds of nucleotides in length from high throughput chip-synthesis. A BASIC code system, a previously developed DNA mediated distributed information storage in our lab, was applied to translate digital binary information to nucleotide base sequence and an encoding redundancy of 1.56% at software level was designed to tolerate the physical dropout of minority oligo. In pushing the limitation, oligo pools comprising of 509 and 11520 distinct oligos, generating the largest population for mixed culture of bacterial cell, were stored. For covering the huge number of oligo population, we assembled them in a redundant fashion and then stored in a mixed culture on solid or liquid medium. Furthermore, the underlying principle of the manufacture of data storage cells was explored with developed deep bioinformatic analysis tools. It demonstrated that oligo homology assembly process is relatively high biased in sequence context and the oligo copy number distribution was more skewed with the assembly fragment number increased. However, after the assembly and transformation, interestingly, it found that the massive oligo remained stable in mixed culture of *E. coli* cells even over multiple passages and remained the quality of digital oligo for perfect information decoding. Finally, it demonstrated that this simple materials of mixed culture of cell achieved in vivo storage of 445 KB digital files in total 2304 Kbps synthesis DNA in a fast and economical way, the largest scale archival data storage in living cell so far, and paved the way for biological data storage taking advantage of both in vitro synthesis capacity and the biological power of living cell in an economical and efficient way, which is crucial for develop practical cold data storage in large scale.

## 2. Results

### 2.1 DNA data storage in mixed cell culture

Thus far, oligo pool comprising of massive distinct oligos are used as material storing archival data in the major in vivo DNA storage approaches. We challenged to merge the advantage of both in vitro oligo pool mediated data storage and in vivo cell system with a novel designed strategy improve the DNA material for data storage. As illustrated in **Figure 1**, binary sequence of archival data was encoded to nucleotide base sequence and spilled into group of oligo strand with few hundreds of nucleotides in length by a BASIC code, which was developed for a DNA oligo pool mediated information distributed storage.^[20]^ In this encoding system, relative low coding redundancy of 1.56% to tolerate the whole oligo physical loss, the dropout. Thus, information could be perfectly decoded as long as more than 98.44% of designed oligo can be retrieved. In addition, oligo strand with letter mutant including base substation or insert/deletion could be corrected by predesigned coding algorithms.^[20]^ Following sequence encoding design, oligo was physically synthesized from the emerging high-throughput chip-based synthesis. Currently, there is only a few commercial products available for massive oligo synthesis, and the quality of oligo pool varied with the manufacture and even the batch. As reported in many previous studies, the unevenness of molecular copy number in oligo pool caused serious problem in the DNA material for data storage.^[21]^ For storing oligo in living cell, oligo could be assembled into high copy number vector plasmid using homology-based cloning method, without any specific sequence, and then the large population of plasmid could be transferred into well-used *E. coli* engineering strain and stored in a mixed culture way. Thus, oligo pool could simply be converted to a living cell-based material for data storage. Mixed culture is a well-used approach majorly in metabolism engineering and direct evolution, which used to generate DNA library with large diversity in living cell. In considering data storage, it requires cell to stably carry these digital DNA sequence in large number. However, there is still short of systematic analysis on how stable the mixed culture carrying large massive oligo will be. Therefore, a multiple-step process, including homology assembly, transformation and mixed culture, was designed to constructed the living cell-based DNA storage. For increasing the homology cloning efficiency, the homology arm sequence was designed with less secondary structure and less cross recognition with each other in NUPACK (Figure S1 and S2).^[22]^ The homology arm was fused with oligo by a PCR amplification through the uniform adapter on both side (**Figure 2**a, Figure S3). In the amplified structure, two Not I cleavage sites were designed on both ends, by which the original oligo sequence could be directed cleaved out from the vector. A redundant assembly is designed to increase the foreign DNA load on each vector. Totally, 6 homology arm sequences were designed for multiple fragments homology assemble into single vector plasmid (Figure S4). Oligos fused with different combination of homology arms could be assembled together. Therefore, in single vector plasmid, 1F, 3F and 5F of fragments could be assembled and each fragment could cover the intact oligo pool. Thus, the multiple fragments assembly principally could largely increase the chance of oligo being assembled into vector plasmid. Following the assembly, circular DNA will be transformed into *E. coli* DH10β cell for mixed culture and then the massive oligos could be retrieved from isolated plasmid.

**Figure 1.**
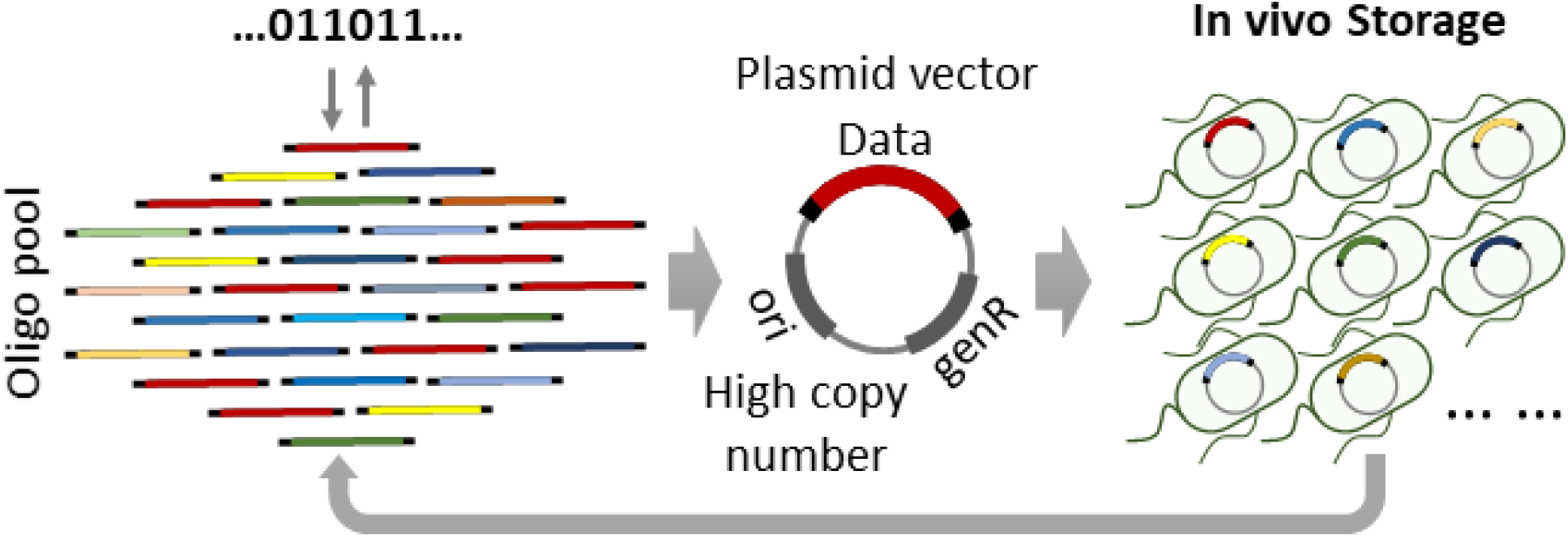
Illustration of mixed culture of bacterial cell for large data storage. First, binary digital information was translated into nucleotide sequence by BASIC encoding system, and then synthesized in a large short oligo pool by chip-based high-throughput synthesis. The oligo pool was assembled into circular plasmid and then transformed into bacterial cell for stable data storage. Oligo pool could be retrieved from the mixed culture of cells for information decoding when need.

**Figure 2.**
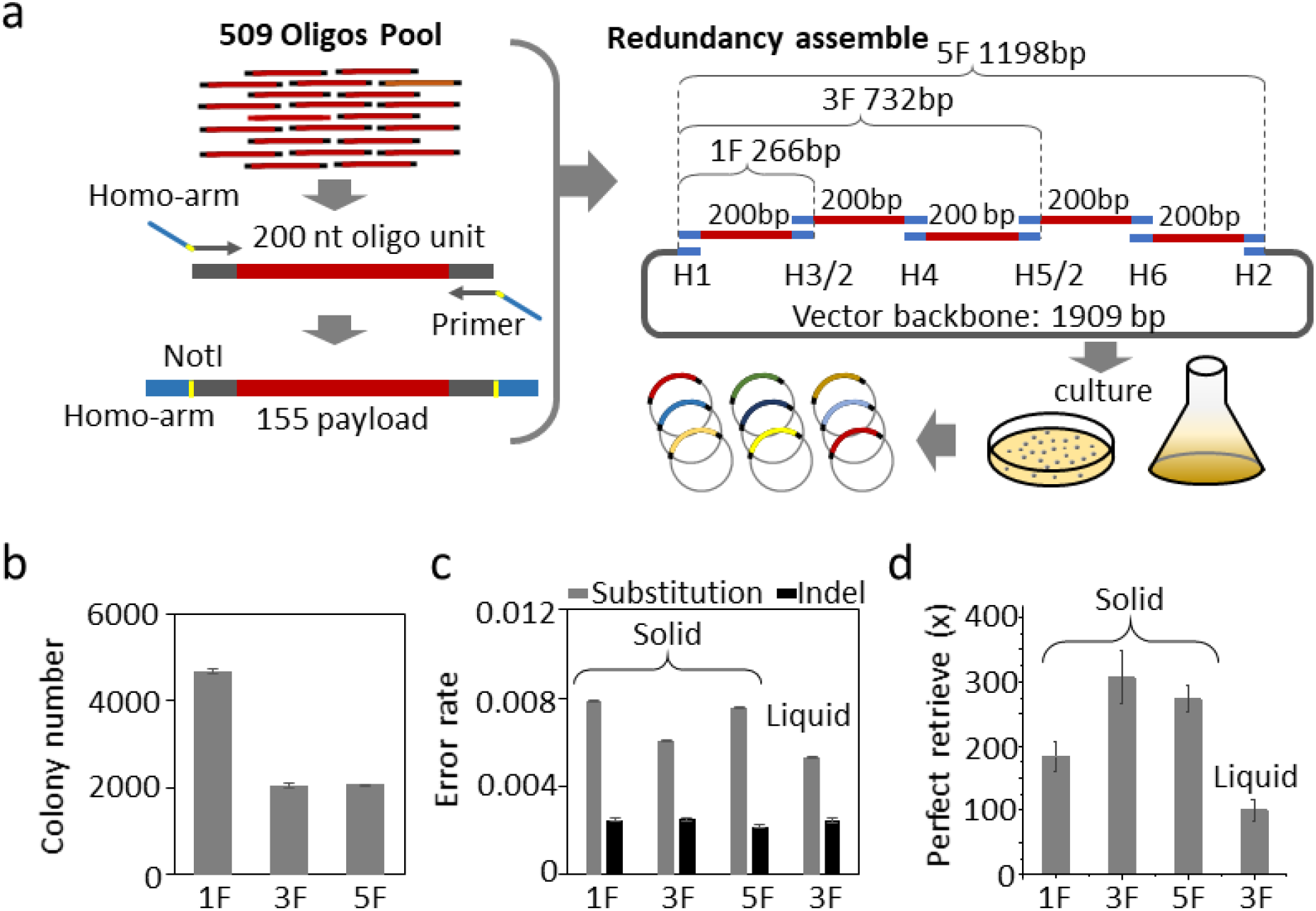
Redundant assembly of 509 oligos pool for mixed culture. a) Schematic for workflow of assembly of DNA pool comprising 509 distinct oligos. Oligos were fused with homology arm via PCR amplification, Not I cleavage site were for oligo retrieve afterwards. Multiple insert fragments, 1F indicate one insert fragment, 3F for three insert fragments and 5F for five insert fragments respectively, each fragment comprising all the 509 oligos, are assembled into a vector plasmid backbone of 1909 bps in length by off-the-shelf homology assembly reagents. Last, the assembled plasmids are transformed in E. coli cell for mixed culture on solid or liquid medium. b) Colony number was counted from solid medium surface for 1F, 3F and 5F assembly. c) Letter error, base substation or indel (both of base insertion or deletion) occurred in oligo pool retrieved from mixed culture on solid or liquid medium and quantified as percentage of counted error base number vs total sequenced base, substation error in gray bar, indel error in dark bar. d) the minimal necessary sequencing reads depth for perfect retrieve of all 509 oligos from 1F, 3F, and 5F assembly sample on solid or liquid medium. Error bars represent the mean ±s.d., where n=3.

### 2.2 Mixed culture of redundant assembled massive oligo pool

Firstly, we tested a pool comprising of 509 distinct oligos as part of a large chip synthesized pool. It known that cell lose its population due to disadvantage in growth rate in mixed culture.^[23]^ With concerning loss of cell carrying minority oligo in the pool, electrically transformed cells were cultured on the surface of solid medium, which should give all cell carrying the assembled plasmid equal change to grow up. The colony number assembled from 1F assembly of total 0.08 pmol oligo fragment and 0.16 pmol vector was counted almost twice of the 3F (assembly of 0.8 pmol each oligo fragment and 0.16 pmol vector) and 5F (assembly of 0.8 pmol each oligo fragment and 0.16 pmol vector) on solid medium surface (Figure 2b and Figure S5-7). There is a trade-off between the assembly efficiency and capability, the redundant assembly could increase the load capability for each vector, but significantly decrease the assembly efficiency. Totally, 122.4 and 158.6 and 268 copy per designed oligo was calculated from the counted colony number for 1F, 3F and 5F respectively. After plasmid isolation, oligo pool was directedly cut out using exonuclease Not I (Figure S8-10) and sequenced by standard NGS. The letter error including substitution or indel were counted, and it was observed that substitution was higher than indel error for all of the assembly samples (Figure 2c) and the error rate is in consistent with previous studies. It was also observed that sequencing reads with single letter error (substitution or indel) was much higher than others (Figure S11), which is in agreement with our previous study as well.^[4]^ For all of the assembly sample, oligo was 100% identified in the sequencing reads, but 1F assembly recorded the low minimal necessary coverage of sequencing reads, at which perfect 100% oligos can be identified (Figure 2d and Figure S12). After the success of oligo retrieve using solid culture, mixed culture in liquid medium was also tested (Figure S13). Plasmid was isolated from 5 ml liquid mixed cell culture and sequenced. The minimal necessary coverage was counted even lower than 1F assembly on solid surface (Figure 2d). Furthermore, the frequency for each oligo counted in the retrieved pool was quantified and similar frequency distribution (Figure S14) were observed for all the assembly samples with very close Gini index (Figure S15). These results demonstrated that the DNA pool of 509 distinct oligos was stably stored in mixed culture.

Next, a DNA pool comprising 11520 distinct oligos with 200 nucleotides in length, over 20 times larger than the first pool, was tested. There is about 445 KB digital files were encoded, including image, word text and virous type files (Figure S3b). It was observed that the mixed culture in liquid medium gave more lower minimal necessary coverage of sequencing reads than solid culture. Additionally, subculture is necessary for long-term storage at low cost. Therefore, the DNA pool with 11520 oligos were assembled to test the subculture of this huge cell population (**Figure 3**a). Totally, the mixed culture was successively passaged 5 times, and plasmid carrying digital DNA were isolated from a large liquid culture and then massive oligos was recovered following Not I digest (Figure S16). There is no obvious difference was observed in the letter error rate even between the 1^st^ and 5^th^ subculture of 1F or 3F assembly samples (Figure 3b). Being in agreement with previous result, the substitution ratio is still higher than indel. From the NGS sequencing reads, some sequences were identified as contamination from host cell genome by deep bioinformatic sequence comparison analysis, but the contamination content is very low, less than 0.2% of the total sequencing reads. This contamination may come from the step of plasmid isolation, because there is also 20 Not I cleavage site on the DH10β genome. But it is very easy to distinguish these contaminations from the true digital oligo sequence by these designed adaptor sequence on the oligo terminal end (Figure S3a). Due to the digital DNA sequence was stored in plasmid, it is still relatively easy to remove the host cell genome contamination clearly in the isolation process just using available commercial bio-reagent. It could be another advantage in comparing with approach, in which digital DNA sequence were directly stored on cell genome.

**Figure 3.**
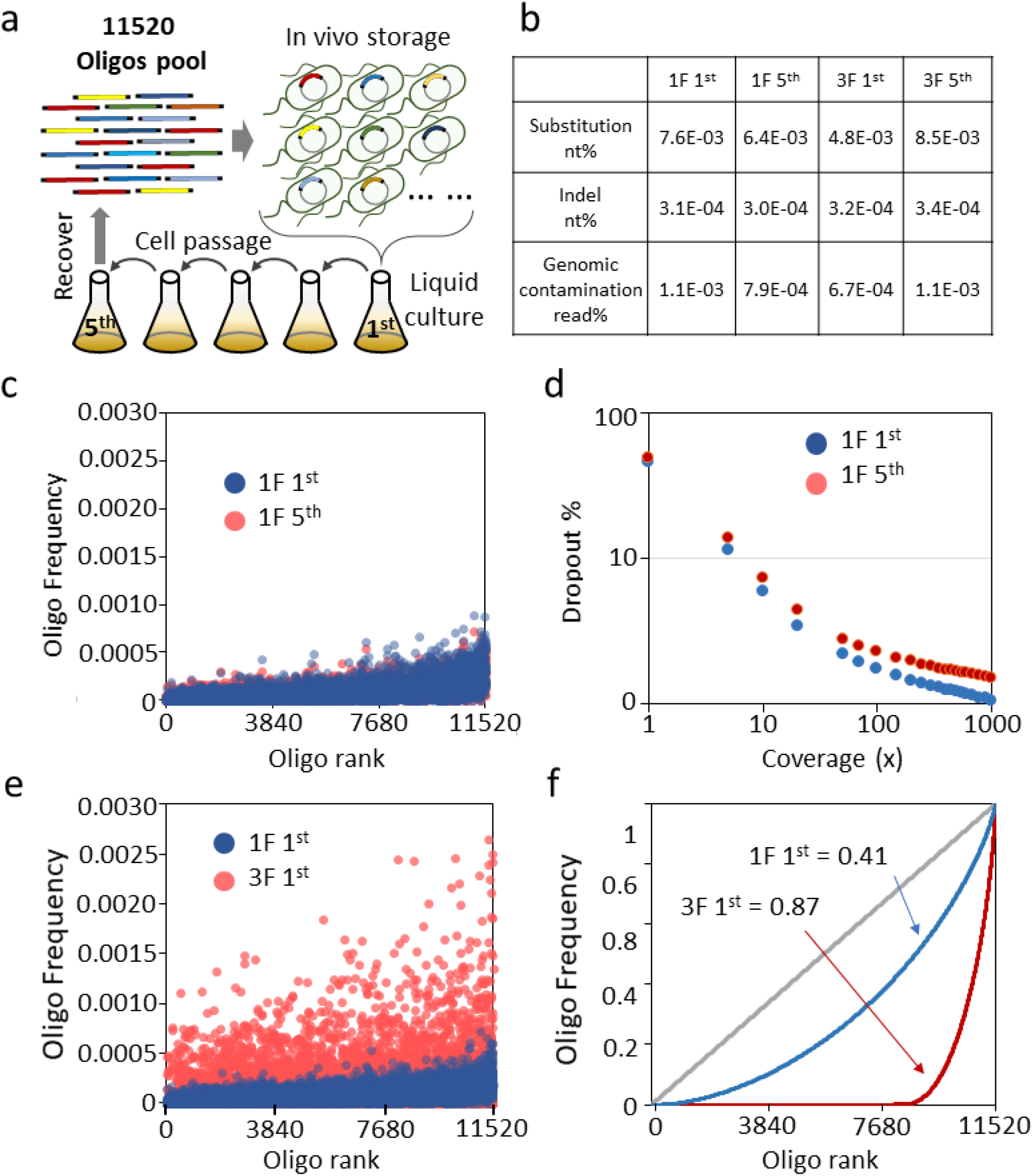
Mixed culture of cells carrying redundant assembled 11520 oligos for large data storage. a) Schematic of cells carrying assembled 11520 oligos pool for successive multiple subculture, cells collected from 1^st^ and 5^th^ passaging were subjected to oligo retrieve and information decoding. b) Letter error rate was quantified form sequenced oligos of 1^st^ and 5^th^ subculture of one insert fragment (1F) or three insert fragments (3F) assembly. The amount of oligo with sequence in high similarity with host cell genome in sequencing reads was identified as genomic contamination. c) The frequency for each of 11520 oligos quantified in sequencing reads from 1^st^ (blue dot) and 5^th^ (red dot) passaging of one fragment (1F) assembly sample. d) Oligo dropout rate was quantified from different sequencing depth (various amount NGS sequencing reads) of 1^st^ (blue dot) and 5^th^ (red dot) passaging of one fragment (1F) assembly sample. e) The frequency for each of 11520 oligos quantified in sequencing reads from the first cell passaging of one insert fragment assembly (1F, blue dot) and three insert fragment assembly (3F, red dot). f) Gini index was quantified for the oligo frequency distribution in the retrieved oligo pool. The 1^st^ passaging of one fragment assembly was quantified as 0.41 (blue line) and 0.87 for 1^st^ passaging of three fragment assembly (red line).

Interestingly, the population of assembled plasmid carrying the inserted digital DNA sequence remained relatively stable. The frequency for each oligo in the pool was not changed significantly in comparing the 1^st^ and 5^th^ passage of 1F or 3F assembly sample (Figure 3c and Figure S17) and the dropout rate decreased when the sequencing going deep (Figure 3d and Figure S18). The bioinformatic analysis demonstrated the stability of oligo pool recovered from the successive passaging. To be surprising, the mixed culture of *E. coli* cells carrying this large population of oligos remained its content uniformity, the Gini index was 0.41 and 0.48 for 1^st^ and 5^th^ of 1F assembly sample respectively (Figure S19). In contrast, the content uniformity was skewed significantly for 3F assembly sample (Figure 3e). In comparing with 1F assembly, in 3F assembly about 21% oligos were enriched accounting for up to 96.2% of the total sequencing reads and the left 79% oligos was largely deprived only accounting for 3.8% of sequencing reads, resulting to a 0.87 of Gini index (Figure 3f). However, the 1^st^ and 5^th^ of 3F assembly sample was relatively consistent with close Gini index and oligo content frequency. The stable oligo frequency distribution even across multiple passaging indicated that the mixed culture of living cell could be qualified materials for data storage.

### 2.3 Large scale DNA data storage in living cell

Thus, living cell mediated DNA data storage was demonstrated in a large scale by a simple multiple-step process, by which DNA pool comprising of massive oligos could be quickly transferred into living cell for data storage (**Figure 4**a). Furthermore, deep bioinformatic analysis explored the underlying principle of this digital storage cell manufacture process. The assembly is found as biased process, its efficiency going down with assembly fragment number increased in the designed redundant assembly. For 11520 DNA pool, much less colony number was counted from 3F assembly than 1F sample and average copy number per designed oligo was calculated as 9.42 for 1F and only 0.91 for 3F assembly sample. Thus, it took more long time for 1^st^ of 3F assembly cell (11 hrs) to reach 1.2 of OD_600_ than 1F assembly cell (8.4 hrs). Even over 1E+6 average molecule copy for each fragment was subjected to the assembly process, but the success assembled copy number for each oligo was quantified only from dozens to hundreds after assembly and transform step. However, the mixed culture amplified the population in a relatively stable fashion without skewing the oligo frequency distribution, probably over 1E+7 average copy of each oligo could be recovered from a batch culture. From these recovered oligos, all of the 1F subculture sample retrieved enough oligo (about 1E+3 copy of each oligo) for perfect information decoding, with finial 0.9% and 1.4% dropout rate for 1^st^ and 5^th^ respectively lower than the 1.56% of decoding limitation. But more oligo was lost in 3F assembly sample, with 26.5% and 32.8% dropout rate for 1^st^ and 5^th^ respectively and similar retrieve rate was obtained in oligo pool recovered by PCR amplification (Figure S20). By mapping the dropout oligo of 1F assembly into the frequency distribution of the original master pool from chip synthesis, it found that the dropout oligo of master pool in sequencing coverage of 10x did not overlap with that of 1F and many oligos in the 1F dropout were mapped to high frequency in master pool (Figure 4b). Furthermore, the enriched oligos group in 3F 1^st^ were also mapped to the frequency distribution of master pool, this group of oligos covered very wide area and mapped to oligos with both high and low coverage (Figure S21). In 10-mer DNA sequence pattern analysis, the top 10% high frequency 10-mer pattern accounted for 42.1% of total 10-mer pattern counts for 3F 1^st^ assembly sample, but the 26.5% for 1F 1^st^ assembly resulting to 16.4% decreasing (Figure S22). The 10-mer frequency distribution was obviously different between the enriched deprived oligos sequence (Figure S23). These results also supported that the assembly process is a biased process dependent on the sequence context rather than the oligo concentration in original master pool. But as long as the living cell materials manufactured, the mixed culture preserved stability of digital DNA for large scale living cell data storage.

**Figure 4.**
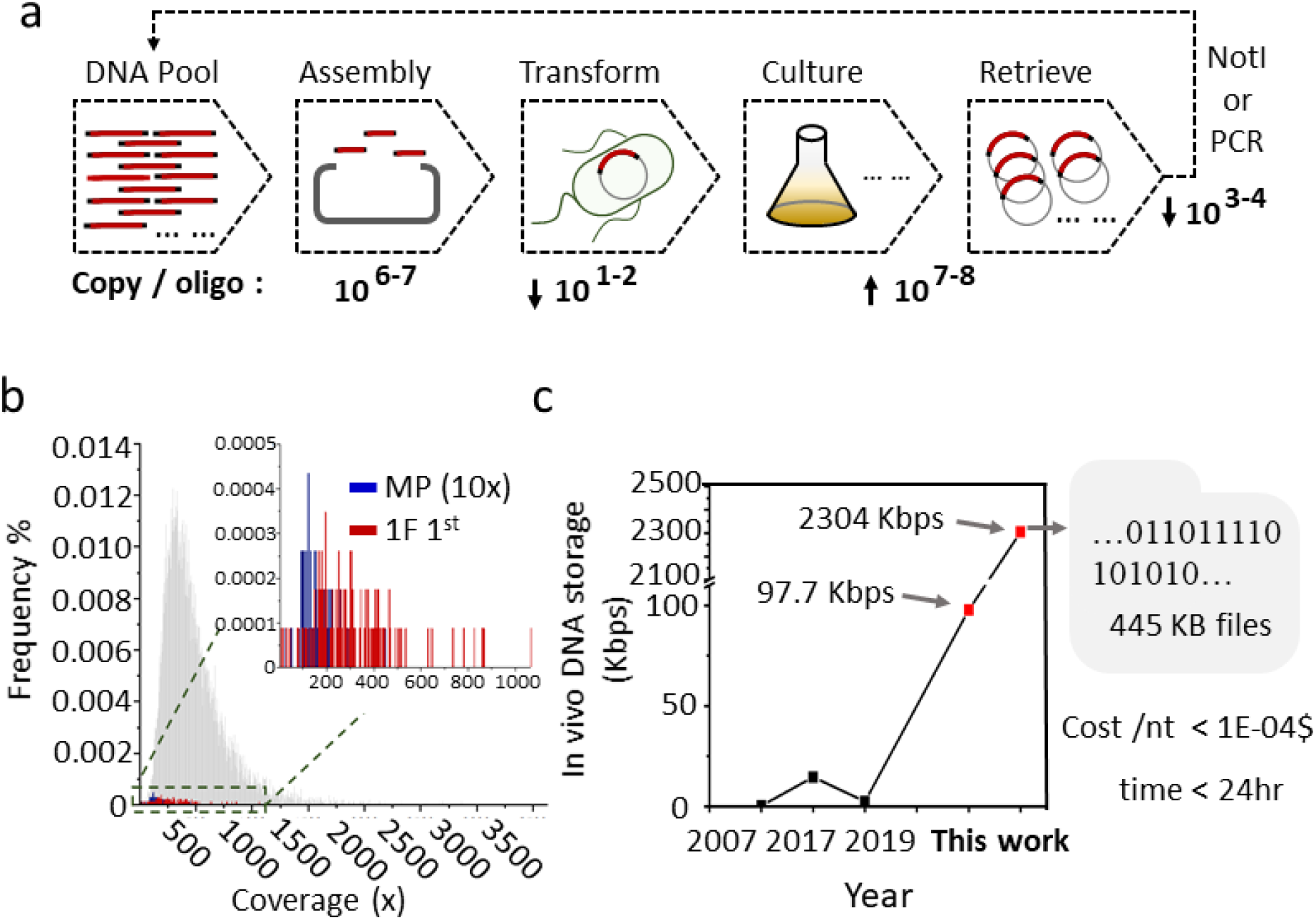
A large-scale DNA data storage in living cell. a) The workflow for the manufacture of mixed culture living cell data storage materials. Oligo pool was assembled with 1E+6⁓7 of average copy of each oligo was subjected to assembly and then transformed into E. coli cell with about 1E+1⁓2 average colony number of each oligo was obtained and then the cell population could be amplified to large scale in mixed culture for further plasmid retrieve and information decoding. b) the 0.9% dropout oligos in 1^st^ passaging of one fragment assembly (red line) and the 0.56% dropout oligos in 10x sequencing reads of original master pool (blue line) were mapped to the oligo frequency distribution of original master pool (gray line). c) In comparison with previous reported major systems for DNA storage in living cell including 0.25 kbps by Yachie in 2007, 18.2 bps by Shipman in 2017 and 2.8 kbps by Sun in 2019, totally 97.7 kbps DNA for 509 oligos pool and 2304 kbps for 11520 oligos pool were stored in mixed culture of E. coli cells at cost lower than 1E-4$ per base and mixed cell storage materials could be manufactured within 24 hrs.

## 3. Discussion

DNA is expected as high-potential material for mass data storage, the serious problem human society will face in the very near future. Beside the storage density, the crucial features including storage longevity and low copy cost are highly dependent on biological system of cell. Thus far, the data storage capability has been demonstrated majorly using massive oligo pool, up to 13 million DNA oligos from the advanced chip synthesis.^[24]^ Although several molecular tools have been adapted from CRISPR and special recombinase to write information into cell genome, the capability is still very far away from in vitro system, not larger than 20K bps so far.^[25]^ Theoretically, one intact single DNA fragment is the desirable material for data storage as the way genome do in nature, but the current DNA writing technology is not designed for long DNA synthesis. Although, the entire bacterial genome has been built up from the chemical synthesized oligos,^[26]^ but large size DNA fragment synthesis requires extreme much labor and time. The cost for DNA fragment over 10 Kbps is about 0.2$/nt at the major commercial company,^[27, 28]^ and generally take over several months to build at high failure risk for complicate sequence. In considering the scale of application, it is hard for large DNA fragment to match for practical data storage until suitable synthesis technology developed. By contrast, oligo pool with several hundreds of nucleotides in length could be synthesized at cost lower than 0.001$/nt,^[27]^ several orders of magnitudes lower than large fragment DNA synthesis, and over million distinct strands could be manufactured at same time in just couple business days and its cost keep going down with synthesis scale going up. Therefore, the mixed culture of bacterial cell carrying massive oligo pool could be a high potential material with advantage of both oligo pool and living cell for data storage. To the best of our knowledge, in comparison with the major previous reported living cell DNA storage system,^[9, 25, 29]^ the total 2304 kbps DNA achieved the largest storage size of data, including text, image documents and computer program code, in living cell (Figure 4c and Supplementary Note 2.7). In comparison with storing long fragment DNA on genome, mixed culture storage materials could be fabricated within 24 hrs after oligo pool synthesis at total manufacture cost, lower than 1E-04$ per base (Supplementary Note 2.3). Thus, in the view of this very artificial approach purpose, digital information storage, it is not necessary to follow the way by which genome information was recorded in nature.

Mixed culture is one technology which has been successfully applied in many fields. In metabolism engineering, different types of microbe cells were cultured together for mutual metabolism benefit,^[30]^ but the size is relatively small. More larger DNA structure with coding huge genomic diversity were generated in living cell for screening of specific biofunction in directed evolution research.^[31]^ Although large DNA library has been created in living cells to generate huge phenotype diversity, but stably carrying these massive DNA structures is not necessary. Generally, it is difficult to balance the growth rate between different cells. In this present work, even in one insert fragment assembly of the massive oligo pool, there is at least 11520 genotype and will be a huge number in the redundant assembly of multiple fragments sample, the largest mixed culture reported so far. However, relative stable mixed culture was achieved even after multiple cell passaging. The copy number distribution of oligos remained stable with very similar value of Gini index in the successive multiple passaged mixed culture (Figure 3d and Figure S15, S19, S24-25). The stability could be considered being supported by a few reasons. The artificial purpose of storing digital information allow designing sequences to avoid sensitive sequence pattern with specific biofunction, e.g., polynucleotides (polyA, polyT, polyC and polyG) and specific exonuclease recognition sequence (Supplementary Note 2.2). The bioinformatic analysis demonstrated that there is no sequence similarity between the designed oligos and the whole *E. coli* DH10β genome with e-value of 1E-6 (Supplementary Note 2.4). It demonstrated that the digital DNA sequence has no significant influence on both host cell growth and the vector plasmid replication. Additionally, storing digital sequence on vector plasmid decreased the information contamination from genome. Therefore, this simple method is highly compatible with any oligo pool for data storage, and scale-up could be achieved easily in a parallel manner based on the over 1E+4 oligo storage we demonstrated here.

In manufacturing of living cell material for data storage, assembly and transformation become crucial step in determining the actual size of oligo population. The deep bioinformatic analysis demonstrated that assembly process is sequence context biased and transformation is a relatively random and inefficient process, the size of oligo population decreased almost two orders of magnitude. The bias occurred in assembly and transformation should highly dependent on the used bioreagent, and homology assembly method should be re-designed to improve its efficiency for assembly of oligo pool with large molecular population. In addition, it found that the dropout rate during mixed culture fit in the dropout curve of master oligo pool, which could be quantified to assess the manufacture of storage material (Figure S26). Therefore, there is still much space to improve the capability of mixed culture cell in storing data. The unevenness of oligo copy number in the original chip-synthesized DNA pool is huge, which is also the serious problem in vitro DNA storage approach.^[21]^ Therefore, more synthetic tools could be developed to improve the chip-synthesized oligo pool and foreign DNA transformation, and balance the large size mixed culture. In summary, DNA oligo pool from chip synthesis comprising of over ten thousand strands was quick transferred into the living cell for data storage, the mixed culture of E. coli cells is a stable material for massive digital DNA sequence and achieved the largest data storage in living cell.

## 4. Experimental Section

### Library construction

For 509 assembly experiment, the oligo pool was synthesized and the lyophilized pool consisted of 11776 oligos of 192 nts, which included the 152 nts payload in each oligo. The pool was resuspended in 1× TE buffer for a final concentration of 2 ng/μL. One of the files, 509 oligos, was flanked by landing sites for primers F01/R01. PCR was performed using Q5® High-Fidelity DNA Polymerases (NEB #M0491) and primers F01/R01 (10 ng oligos, 2.5 μL of each primer (100 mM), 0.5 μL Q5 High-Fidelity DNA Polymerases, 4 μL 2.5 mM dNTPs in a 50 μL reaction). Thermocycling conditions were as follows: 5 min at 98 °C; 10 cycles of: 10 s at 98 °C, 30 s at 56 °C, 30 s at 72 °C, followed by a 5 min extension at 72 °C. The library was then purified using Plus DNA Clean/Extraction Kit (GMbiolab Co, Ltd. #DP034P) and eluted in 40 μL ddH2O. This library was considered the master pool and run on the 2% agarose gel to verify the correct size. For 11520 assembly experiment, the synthetic DNA pool consisted of 11520 oligos of 200 nts, which included the 155 nts payload flanked by landing sites for primers F02/R02 (Figure S3). The lyophilized pool was rehydrated in 1× TE buffer and used the above protocol to amplify the file.

### DNA storage in living cells

For the 509 oligos pool assembly fragment preparation, we started with the master pool as described above. The fragments were prepared with different homologous arms using Q5® High-Fidelity DNA Polymerases and the corresponding primers. Then the Gibson Assembly® Master Mix – Assembly (NEB, #E2611) was used according to user’s manual. For the 11520 oligos pool assembly fragment preparation, we started with the master pool as described above. The fragments were prepared with different homologous arms using 2×EasyTaq® PCR SuperMix (AS111, TRANS) and the corresponding primers. NEBuilder® HiFi DNA Assembly Cloning Kit (NEB, #E5520) was used according to user’s manual. After assembly, the constructed samples were transformed into DH10β electrocompetent cells. The information about experimental procedures was detailed in supporting information.

### Data recovery

After liquid and plate culture, the plasmid was extracted using plasmid minipreparation Kit (TIANGEN, #DP103), respectively. Then QuickCut™ Not I (Takara, #1623) was used for fragments recovery. After gel cut by Plus DNA Clean/Extraction Kit, the samples of 509 oligos pool (1 F, 3 F and 5 F) and 11520 oligos (passage-1 and passage-5 of 1F and 3 F) were sequenced directly. To get more complete information, we performed a PCR amplify process from constructed plasmid to amplify 11520 oligos (passage-1 and passage-5 of 1F and 3 F) using Q5® High-Fidelity DNA Polymerases and primer set F02/R02. The thermocycling protocol was: (1) 98 °C for 5 min, (2) 98 °C for 30 s, (3)54 °C for 30 s, (4) 72 °C for 10 s, then repeat steps 2–4 five times. Finally, the PCR reaction was terminated at 72 °C for 5 min, and purified using Plus DNA Clean/Extraction Kit (GMbiolab Co, Ltd. #DP034P) then sequenced them.

## Supporting Information

Supporting Information is available from the Wiley Online Library or from the author.

## Acknowledgements

This work was supported by National Science Foundation of China (Grant No.21476167, No.21778039 and No.21621004). M. H., Y. G. and H. Qiao contributed equally to this work.

