## Supplemental file for "Mixed Culture of Bacterial Cell for Large Scale DNA Storage"

Dr. Xin Chen

Center for Applied Mathematics, Tianjin University, Tianjin 300000, China.

Key Laboratory of Systems Bioengineering (Ministry of Education), School of Chemical Engineering and Technology, Tianjin University, Tianjin 300000, China.

† Equal contribution

### 1. Supporting Materials and Methods

#### 1.1. Strains and Culture conditions

Electrocompetent competent *E. coli* DH10β was used for cloning and purchased from Biomed Co., Ltd. (Beijing, China). The antibiotics of ampicillin were used at 100 mg/mL. Unless otherwise noted, the cells were cultured in Luria-Bertani (LB) broth with shaking at 220 rpm or on LB agar plates at 37 °C.

#### 1.2 Computational strategy for primer homologous arm design

The primers were included three parts, one was the homologous arm (Homo-arm) which used for gibson assembly, the other was Not I which endonuclease recognition site and the last was address sequence. The primer design algorithm was implemented to codify rules for assembly fragments production using PCR. Primer homologous arm was designed using the NUPACK (http://www.nupack.org). Sets of primers are identified satisfying the following rules for homologous arm design: (1) homologous arm length 25 nt; (2) guanine-cytosine (GC) content between 40 and 60%; (3) no interaction between each homologous arm. Selected primers passing all of the aforementioned tests are provided as output as correct pairs that would yield the product of the defined size (Figure. S1 and S2).

#### 1.3 Assembly experiment

For the backbone preparation, the pUC19 plasmid was used as templates for PCR. The PCR was performed using PrimeSTAR® Max DNA Polymerase (Takara #R045Q) and primer set PCR-vactor-F/R. Thermocycling were amplified for 30 cycles, involving denaturation for 15 s at 98 °C, annealing for 5 s at 55 °C and primer extension for 20 s at

72 °C. Then the PCR products were purified by gel cut using Plus DNA Clean/Extraction Kit (GMbiolab Co, Ltd. #DP034P) and eluted in 30 μL dd$H_2O$.

For the 509 oligos pool assembly fragment preparation, we started with the master pool as described above. The fragments were prepared with different homologous arms using Q5® High-Fidelity DNA Polymerases and the corresponding primers (the sequence was showed in Table S1). The template used 0.2 ng of oligo input from the master pool into 50 μL PCR reactions. Initial denaturation was carried out at 98 °C for 5 min. Follow this by 20 PCR cycles, involving denaturation for 30 s at 98 °C, annealing for 30 s at 56 °C and primer extension for 20 s at 72 °C. Finally, the PCR reaction was terminated by incubating the solution at 72 °C for 5 min. The library was then purified using Plus DNA Clean/Extraction Kit (Gmbiolab, #DP034P) and eluted in 30 μL dd$H_2O$.The final library was run under the same conditions as described. Then the Gibson Assembly® Master Mix – Assembly (NEB, #E2611) was used according to user's manual.

For the 11520 oligos pool assembly fragment preparation, we started with the master pool as described above. The fragments were prepared with different homologous arms using 2×EasyTaq® PCR SuperMix (AS111, TRANS) and the corresponding primers (the sequence was showed in Table S1 and Figure S3). The template used 20 ng of oligo input from the master pool into 50 μL PCR reactions. Initial denaturation was carried out at 94 °C for 2 min. Follow this by 10 PCR cycles, involving denaturation for 30 s at 94 °C, annealing for 30 s at 53 °C and primer extension for 20 s at 72 °C. Finally, the PCR reaction was terminated by incubating the solution at 72 °C for 5 min. The library was then purified and eluted in 100 μL dd$H_2O$. NEBuilder® HiFi DNA Assembly Cloning Kit

(NEB, #E5520) was used according to user's manual.

The concentration of each fragment was calculated for optimal assembly, based on fragment length and weight, we used the following equation:

Equation 1. pmols = (weight in ng) × 1,000 / (base pairs × 650 daltons).

Based on this calculation, the number of molecules per oligo and copy number was determined according to the following formula:

Equation 2. moles dsDNA (mol) = mass of dsDNA (g) / ((length of dsDNA (bp) × 607.4) + 157.9 g/mol;

Equation 3. DNA copy number = moles of dsDNA × 6.022e23 molecules/mol.

We found an excellent agreement concentration between the assembly fragment and the backbone of Gibson assembly experiment. Thus, for 509 oligos pool assembly experiment, optimized cloning was performed using $10^{11}$ copy number of vectors and $10^{8}$ copy number of inserts. For 11520 oligos pool assembly experiment, optimized cloning was performed using $10^{10}$ copy number of vectors and $10^{10}$ copy number of inserts. The sample was incubated in a thermocycler at 50 °C for 60 min, respectively. Following incubation, store samples on ice or at -20 °C for subsequent transformation.

**1.4 Transformation and culture**

Electroporation was carried out in 1-mm gap cuvettes with conditions 1.8 kV, 200 Ω, 25 Mf, cells were recovered into fresh SOB medium for 1 h at 37 °C. For 509 assembly experiment, each sample (5 μL) was added to DH10β electrocompetent cells (50 μL) for electroporation reaction. After recovery, 500 μL cells were plated on the selection medium plates with appropriate selective condition (Amp) and the colonies were counted by

ImageJ. For 11520 assembly experiment, each sample (2 μL, totally 20 μL) was added to DH10β electrocompetent cells (50 μL) for electroporation reaction. After recovery, 500 μL cells were plated on the selection medium plates with appropriate selective condition (Amp) and the colonies were counted by ImageJ. The transformation rate was calculated by the following equation:

Equation 4. Transformation efficiency (cfu/μg) = colonies on plate/ plasmid DNA spread on plate

Equation 5. Transformation rate = Transformation efficiency/ $10^{10}$ (cfu/μg)*

*note: $10^{10}$ cfu/μg is the theoretical transformation efficiency of DH10β electrotransducer cells.

For 509 oligos pool assembly experiment, another 500 μL cells were inoculated in 5 mL Luria broth (LB) medium plus appropriate antibiotics and grown overnight [37 °C, 220 revolutions per minute (RPM)] to obtain seed cultures. For 11520 oligos pool assembly experiment, 5 mL recovered cells were inoculated in 45 mL Luria broth (LB) medium plus appropriate antibiotics and grown overnight to obtain seed cultures. The seed cultures were then serially diluted (1:10) in 50 ml of prewarmed LB plus appropriate antibiotics followed by $OD_{600}$ reached 1.2. This consecutive procedure was repeated 5 times (Figure S12).

**Quantitative and statistical analysis**

Histograms were generated using Origin software. Quantitative data in figures are presented as the mean and standard deviation from three biological replicates.

#### 2. Supplementary Note

##### 2.1 BASIC Code

Gene coding is a new type of distributed storage system. In this work, we use the BASIC Code, it is a kind of distributed erasure code designed for gene coding, aiming at maximizing storage utilization, effectively guaranteeing the reliability of the storage system. Due to the adjustable system parameters K and L, we take the standard system parameters (K =252, L =256) as an example. Here, we use 11,520 DNA sequences (12K oligo pool) of length 200 nt with payload of length 155 nt to store 445 KB data (Figure S3).

##### 2.2 Encoding and Decoding Process

**Encoding process.** The goal of this work is to transform the input file to DNA sequence reads (within biochemical constraints). DNA BASIC Code should enable error-detection, error-correction and full recovery. There are mainly steps: (a) erasure coding, (b) RS coding. (c) filtering. Since the sequence reads need to satisfy the biochemical constraints, both (a) and (b) include the step of filtering the sequences.

**Decoding process.** The decoding process is processed step by step in reverse by the encoding process. XOR processing is performed according to the mapping table to restore the RS code, and then the RS code is used for error correction to ensure that existing information of each sequence is accurate. Restore the BASIC code sequence. For each group of data information, decode according to the BASIC decoding algorithm.

##### 2.3 Cost calculation

In this study, the cost of practical implementation of DNA storage in vivo was $0.001 per base, which was consisted of four parts: DNA synthesis, the DNA library was synthesized from Twist Bioscience and CustomArray, and the cost about $ 0.0009 per base; Assembly, this part was contained PCR and assembly, the cost around $ 58 during an experiment; Transformation, the cost of 509 oligos and 11520 oligos was $ 9 and $ 90, respectively; Recovery, which was mainly included plasmid extraction, enzyme digestion and sequencing.

##### 2.4 The bioinformatic statistical analysis.

We stitched the reads pair by using PEAR to get the sequenced reads.

The sequenced reads were aligned with the actual sequences (synthesized by Twist Bioscience and CustomArray) by basic sequence alignment program (BLAST). The coverage and number could be achieved by Valid_Coverage_Number.pl. The frequency was calculated via the number dividing by total number of actual sequences. Then the distribution of number of reads per each actual sequence was displayed (Figure S24, S25). Valid DNA sequence named payload obtained by Obtain_Payload.pl and kmer of these payload sequences were analyzed by kmer.pl (Figure S22, S23).

The number of each sequence of the sequenced reads could be achieved by Valid_Coverage_Number.pl. The oligo frequency was obtained through the number of each sequence dividing by the sum of these numbers and the distribution of oligo frequency could be displayed (Figure 3c, 3e, Figure S14, S17). Gini index was calculated by R (Figure 3f, Figure S15, S19). Then the frequency was obtained through the oligo frequency of each sequence in 3F 1$^{st}$ dividing by the corresponding frequency in 1F 1$^{st}$, and then counted the distribution of sequences with increased frequency (the frequency was over 1) in oligo master pool (Figure S21).

##### 2.5 Genome blast

**Genomic contamination rate (%).** The sequences with high similarity to the actual sequences were removed by unmatch_test.pl. The remaining sequences were aligned with the genomic sequences of DH10β (from the competent cell used in this work) by BLAST to obtain the unmatched sequence reads (namely genomic contamination reads), and genomic contamination rate was calculated via the unmatched sequence reads dividing by the total sequenced reads (Genomic contamination reads% of Figure 3b). In this work, the default threshold set at blast is e-value $10^{-6}$. The smaller the e-value, the higher the similarity according to NCBI. At the same time, the actual sequences were also aligned with the genomic sequences by BLAST (e-value $10^{-6}$), but there was no output. Then the actual sequences were aligned with the genomic sequences by blast on NCBI, the output result was: No significant similarity found.

##### 2.6 Calculating error rate and dropout (%).

All sequenced reads were aligned with the actual reference sequences by BLAST to screen out the reads with errors containing substitution, insertion, and deletion (hereinafter referred to as "errors") on the payload of a single sequence. The number of reads with an error, two errors, three errors, ……, ten errors, more than ten errors in individual sequences were counted in detail by Mismatch_Analysis.pl and Gap_Analysis.pl, and the frequency were calculated through the number of these reads dividing by the total number of noisy reads (Figure 2c, Figure 3b, Figure S11).

Dropout was calculated by Vaild_Coverage_Number.pl, Random_Access.pl and Dropout.pl (Figure 2d, Figure 3d, Figure S12-S13, S18, S20, S26). According to our encoding strategy which allows a maximum of 4 DNA sequences to be lost or corrupted in each group (each group contains 256 DNA sequences), the redundancy can be calculated ideally as: 4/256=1.56%.

##### 2.7 Calculating of storage size

The data of Figure 3c was calculated from corresponding references, the detail was shown as follow: In 2007, Nozomu Yachie et al. has been inserted redundantly oligonucleotide (C1, C2, C3 and C4) into multiple loci of the *Bacillus subtilis* genome. The size of C1was 64 nt, C2, C3 and C4 were 62 nt, respectively. We calculated the total base is 0.25 Kbps.; In 2017, Seth L Shipman et al. has been encoded images and a short movie into the genomes of a population of living bacteria. The short movie was encoded by five frames of Eadweard Muybridge’s Horse in Motion. Each Frame was represented by a unique oligo set of 104 protospacers, and each protospacer included 28 bases. We calculated the total base is 14.56 Kbps. In 2019, Jian Sun et al. has been stored a poem of “Snow” in *E. coli*, *yeast* and *Arabidopsis*. The sequence of was “Snow” was 2448 base (2.448 Kbp).

#### 3. Supplementary Table

**Table S1**. Sequence information of primers

| Primer | Sequence |
|---|---|
| F01 | ACCCTCACCTATCAACTCAA |
| R01 | CTTCCGACCACTATACCTCT |
| F02 | TGCATCACCTACCTCAGC |
| R02 | TCCACGACGATCAGACT |
| 1-F-1 | TATCCCCTGATTCTGTGGATAACCGGCGGCCGCTCACCATCCACTCTAAACAC |
| 1-F-2 | CTTGGGTTTGTTGGGTTTGTTAGGTGCGGCCGCTCACCATCCACTCTAAACAC |
| 1-F-3 | ACAGAGTAAAGCAAGACCGGATAACGCGGCCGCTCACCATCCACTCTAAACAC |
| 1-F-4 | TTGTAGGTTGGGTGGGTGTGTGGGAGCGGCCGCTCACCATCCACTCTAAACAC |
| 1-F-5 | CAAGCGGTGAGTAATGGAAATCTATGCGGCCGCTCACCATCCACTCTAAACAC |
| 1-R-1 | ACCTAACAAACCCAACAAACCCAAGGCGGCCGCCACTTTACACCTCCACTCAT |
| 1-R-2 | GTTATCCGGTCTTGCTTTACTCTGTGCGGCCGCCACTTTACACCTCCACTCAT |
| 1-R-3 | TCCCACACACCCACCCAACCTACAAGCGGCCGCCACTTTACACCTCCACTCAT |
| 1-R-4 | ATAGATTTCCATTACTCACCGCTTGGCGGCCGCCACTTTACACCTCCACTCAT |
| 1-R-5 | TATAAAAATAGGCGTATCACGAGGCGCGGCCGCCACTTTACACCTCCACTCAT |
| 2-F-1 | TATCCCCTGATTCTGTGGATAACCGGCGGCCGCACCCTCACCTATCAACTCAA |
| 2-F-2 | ACCTAACAAACCCAACAAACCCAAGGCGGCCGCCTTCCGACCACTATACCTCT |
| 2-F-3 | CTTGGGTTTGTTGGGTTTGTTAGGTGCGGCCGCACCCTCACCTATCAACTCAA |
| 2-F-4 | GTTATCCGGTCTTGCTTTACTCTGTGCGGCCGCCTTCCGACCACTATACCTCT |
| 2-F-5 | ACAGAGTAAAGCAAGACCGGATAACGCGGCCGCACCCTCACCTATCAACTCAA |
| 2-R-1 | TCCCACACACCCACCCAACCTACAAGCGGCCGCCTTCCGACCACTATACCTCT |
| 2-R-2 | TTGTAGGTTGGGTGGGTGTGTGGGAGCGGCCGCACCCTCACCTATCAACTCAA |
| 2-R-3 | ATAGATTTCCATTACTCACCGCTTGGCGGCCGCCTTCCGACCACTATACCTCT |
| 2-R-4 | CAAGCGGTGAGTAATGGAAATCTATGCGGCCGCACCCTCACCTATCAACTCAA |
| 2-R-5 | TATAAAAATAGGCGTATCACGAGGCGCGGCCGCCTTCCGACCACTATACCTCT |
| 3-F-1 | TATCCCCTGATTCTGTGGATAACCGGCGGCCGCACTCCCACTCACCTATATCC |
| 3-F-2 | CTTGGGTTTGTTGGGTTTGTTAGGTGCGGCCGCACTCCCACTCACCTATATCC |
| 3-F-3 | ACAGAGTAAAGCAAGACCGGATAACGCGGCCGCACTCCCACTCACCTATATCC |
| 3-F-4 | TTGTAGGTTGGGTGGGTGTGTGGGAGCGGCCGCACTCCCACTCACCTATATCC |
| 3-F-5 | CAAGCGGTGAGTAATGGAAATCTATGCGGCCGCACTCCCACTCACCTATATCC |
| 3-R-1 | ACCTAACAAACCCAACAAACCCAAGGCGGCCGCATAACCTCACTCACCTACCA |
| 3-R-2 | GTTATCCGGTCTTGCTTTACTCTGTGCGGCCGCATAACCTCACTCACCTACCA |
| 3-R-3 | TCCCACACACCCACCCAACCTACAAGCGGCCGCATAACCTCACTCACCTACCA |
| 3-R-4 | ATAGATTTCCATTACTCACCGCTTGGCGGCCGCATAACCTCACTCACCTACCA |
| 3-R-5 | TATAAAAATAGGCGTATCACGAGGCGCGGCCGCATAACCTCACTCACCTACCA |
| 4-F-1 | TATCCCCTGATTCTGTGGATAACCGGCGGCCGCACTCTCACCTTTACTCCCAC |
| 4-F-2 | CTTGGGTTTGTTGGGTTTGTTAGGTGCGGCCGCACTCTCACCTTTACTCCCAC |
| 4-F-3 | ACAGAGTAAAGCAAGACCGGATAACGCGGCCGCACTCTCACCTTTACTCCCAC |
| 4-F-4 | TTGTAGGTTGGGTGGGTGTGTGGGAGCGGCCGCACTCTCACCTTTACTCCCAC |
| 4-F-5 | CAAGCGGTGAGTAATGGAAATCTATGCGGCCGCACTCTCACCTTTACTCCCAC |

| | |
|---|---|
| 4-R-1 | ACCTAACAAACCCAACAAACCCAAGGCGGCCGCCTACTCCCACTACTACCACA |
| 4-R-2 | GTTATCCGGTCTTGCTTTACTCTGTGCGGCCGCCTACTCCCACTACTACCACA |
| 4-R-3 | TCCCACACACCCACCCAACCTACAAGCGGCCGCCTACTCCCACTACTACCACA |
| 4-R-4 | ATAGATTTCCATTACTCACCGCTTGGCGGCCGCCTACTCCCACTACTACCACA |
| 4-R-5 | TATAAAAATAGGCGTATCACGAGGCGCGGCCGCCTACTCCCACTACTACCACA |
| PCR-pUC19-F | GCCTCGTGATACGCCTATT |
| PCR-PUC19-R | CGGTTATCCACAGAATCAGG |
| TY-primer 1-F | TATCCCCTGATTCTGTGGATAACCGGCGGCCGCATCACCTACCTCAGCTC |
| TY-primer 1-R | ACCTAACAAACCCAACAAACCCAAGGCGGCCGCTCCACGACGATCAGACT |
| TY-primer 2-F | CTTGGGTTTGTTGGGTTTGTTAGGTGCGGCCGCATCACCTACCTCAGCTC |
| TY-primer 2-R | GTTATCCGGTCTTGCTTTACTCTGTGCGGCCGCTCCACGACGATCAGACT |
| TY-primer 3-F | ACAGAGTAAAGCAAGACCGGATAACGCGGCCGCATCACCTACCTCAGCTC |
| TY-primer 3-R | TCCCACACACCCACCCAACCTACAAGCGGCCGCTCCACGACGATCAGACT |
| TY-primer 4-F | TTGTAGGTTGGGTGGGTGTGTGGGAGCGGCCGCATCACCTACCTCAGCTC |
| TY-primer 4-R | ATAGATTTCCATTACTCACCGCTTGGCGGCCGCTCCACGACGATCAGACT |
| TY-primer 5-F | CAAGCGGTGAGTAATGGAAATCTATGCGGCCGCATCACCTACCTCAGCTC |
| TY-primer 5-R | TATAAAAATAGGCGTATCACGAGGCGCGGCCGCTCCACGACGATCAGACT |

#### 4. Supplementary Figures

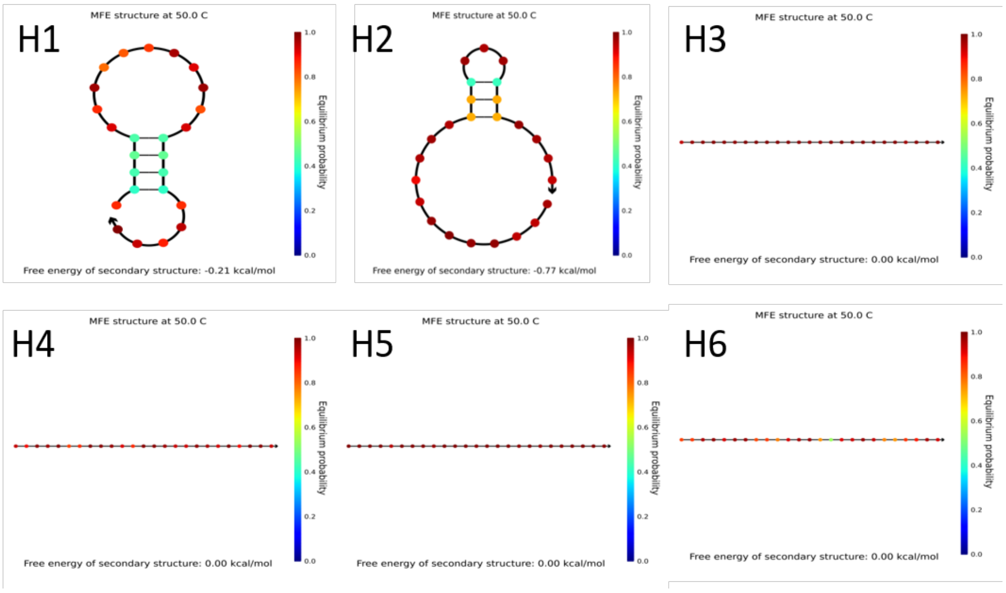


**Figure S1.** Thermodynamic secondary structure in designed homologous arm sequence calculated by NUPACK (http://www.nupack.org). H1, H2 is the homologous arm with sequence from vector plasmid pUC19; H3, H4, H5 and H6, are the in silico designed sequence for homologous assembly.

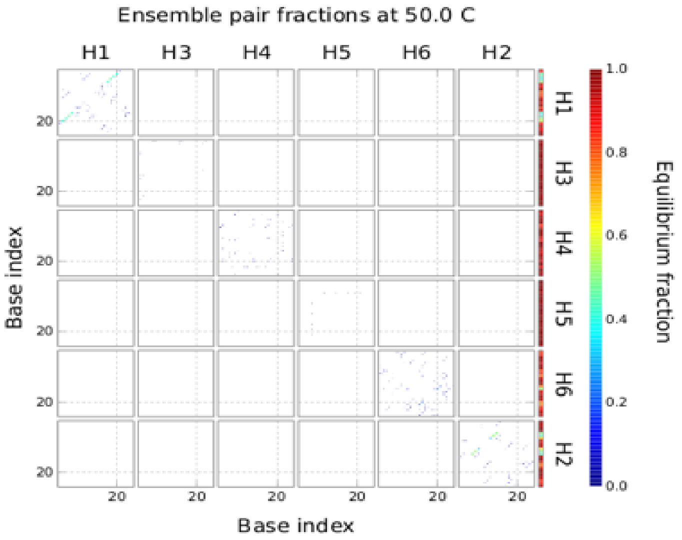


**Figure S2.** Thermodynamic diagram for cross interaction between designed homologous arm sequence assessed by NUPACK. H1, the left homologous arm on the pUC19; H2, the right homologous arm on the pUC19; H3, H4, H5 and H6, the designed-homologous arms.

a

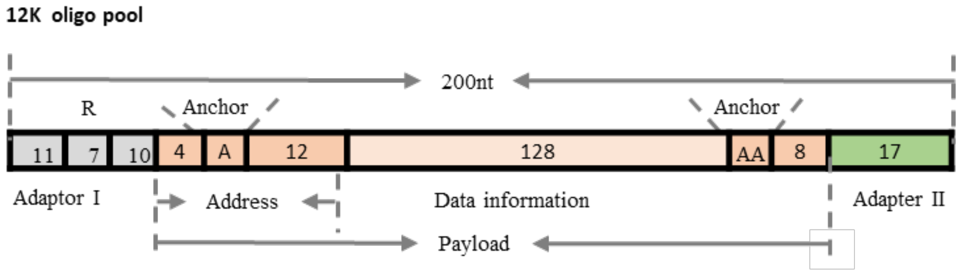


b

| Data | File Size | Number of DNA strands |
|---|---|---|
| Central dogma (.jpg) | 35 KB | 990 |
| DNA helix (.gif) | 81 KB | 2292 |
| China Classical literature (.txt) | 164 KB | 4641 |
| A Brief History of Element (.txt) | 34 KB | 962 |
| Panda burn incense (.rar) | 66 KB | 1867 |
| Human Mitochondrial | 65 KB | 768 |
| Total | 445KB | 11520 |

**Figure S3.** The information of 11520 oligos pool. (a) Structure of the oligos unit designed under the same principle of our previous study, and synthesized from chip-based 12k oligo synthesis product of Twist Bio. (b) The list of 445 KB digital files encoded in the 11520 oligos pool.

| primer | Sequence | | |
|---|---|---|---|
| | Homologous arm | NcoI | Primer |
| TY-primer 1-F | TATCCCCTGATTCTGTGGATAACCG | GCGGCCGC | ATCACCTACCTCAGCTC |
| TY-primer 1-R | ACCTAACAAACCCAACAAACCCAAG | GCGGCCGC | TCCACGACGATCAGACT |
| TY-primer 2-F | CTTGGGTTTGTTGGGTTTGTTAGGT | GCGGCCGC | ATCACCTACCTCAGCTC |
| TY-primer 2-R | GTTATCCGGTCTTGCTTTACTCTGT | GCGGCCGC | TCCACGACGATCAGACT |
| TY-primer 3-F | ACAGAGTAAAGCAAGACCGGATAAC | GCGGCCGC | ATCACCTACCTCAGCTC |
| TY-primer 3-R | TCCCACACACCCACCCAACCTACAA | GCGGCCGC | TCCACGACGATCAGACT |
| TY-primer 4-F | TTGTAGGTTGGGTGGGTGTGTGGGA | GCGGCCGC | ATCACCTACCTCAGCTC |
| TY-primer 4-R | ATAGATTTCCATTACTCACCGCTTG | GCGGCCGC | TCCACGACGATCAGACT |
| TY-primer 5-F | CAAGCGGTGAGTAATGGAAATCTAT | GCGGCCGC | ATCACCTACCTCAGCTC |
| TY-primer 5-R | TATAAAAATAGGCGTATCACGAGGC | GCGGCCGC | TCCACGACGATCAGACT |

**Figure S4.** The primer sequence for 11520 oligos pool insert fragment construction. Homologous arm sequence was indicted in black, Not I cleavage site in blue and primer sequence in red (forward) and green (reverse).

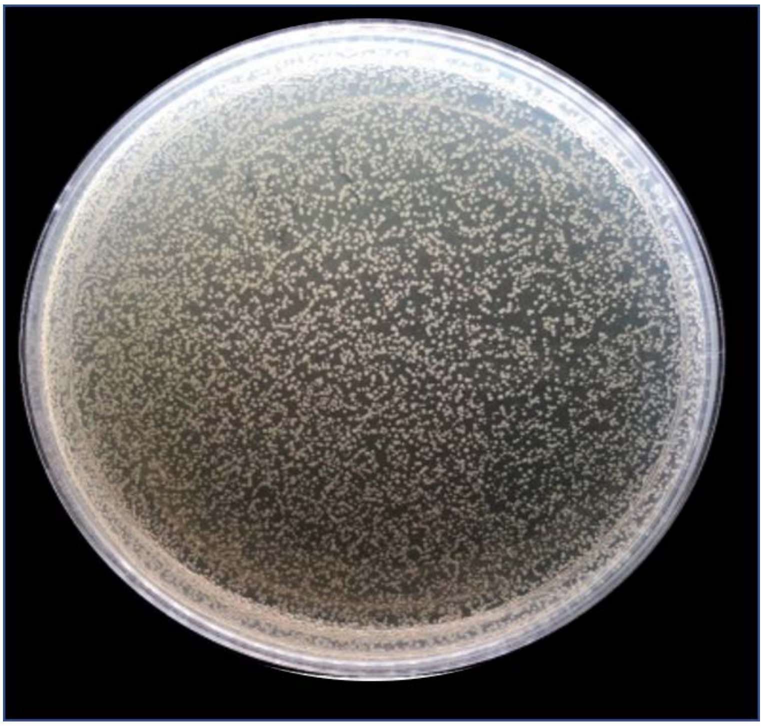

**Figure S5.** One petri-dish of solid medium for one insert fragment (1F) assembly of 509 oligos pool, the colony number was counted from all the solid medium plates.

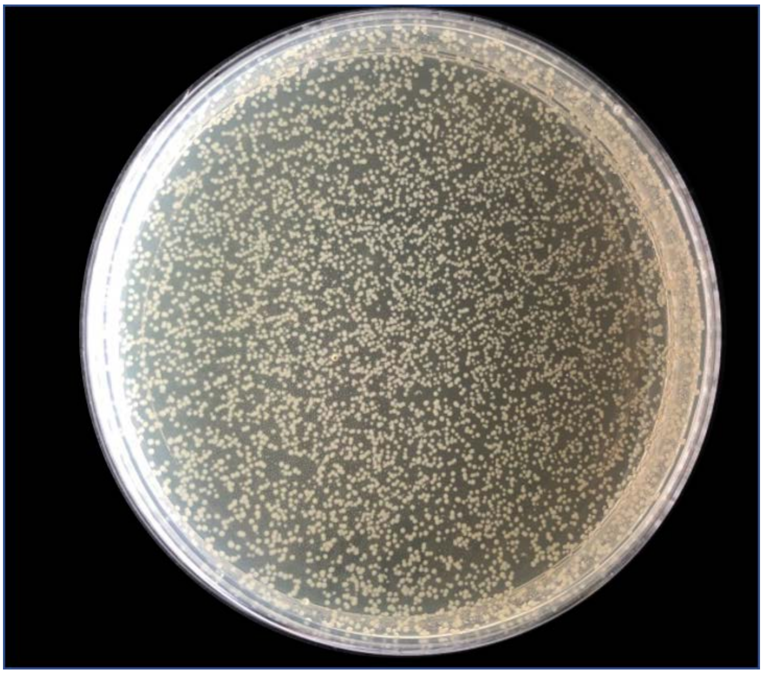

**Figure S6.** One petri-dish of solid medium for one insert fragment (3F) assembly of 509 oligos pool, the colony number was counted from all the solid medium plates.

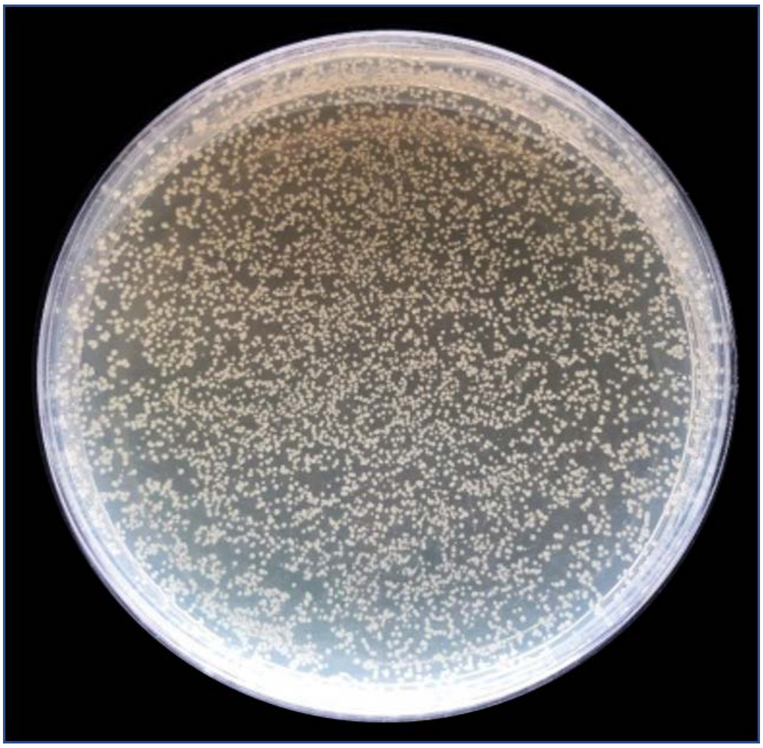

**Figure S7.** One petri-dish of solid medium for one insert fragment (5F) assembly of 509 oligos pool, the colony number was counted from all the solid medium plates.

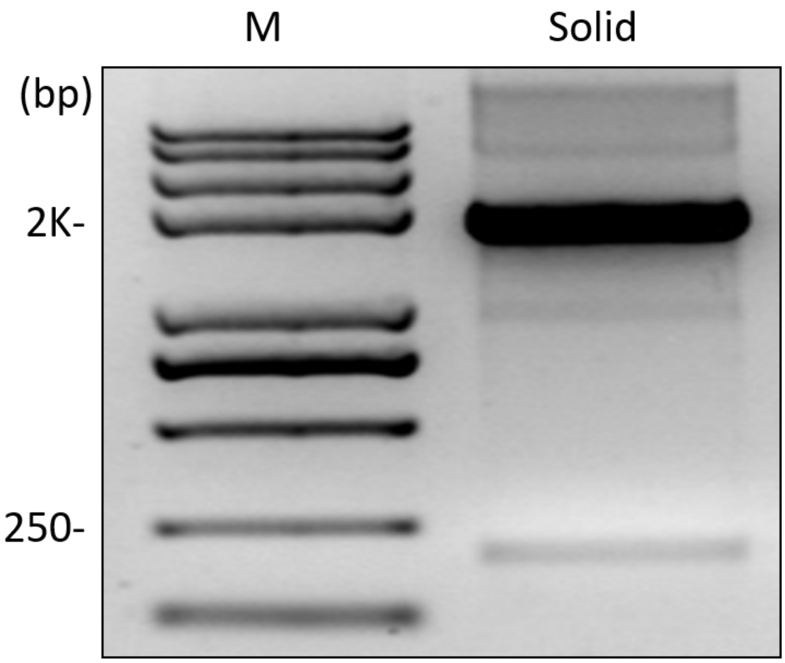


**Figure S8.** Digestion of 1F assembly plasmids by Not I (509 oligos pool). M, 2K plus II DNA marker; Solid, cultured on LB plate medium.

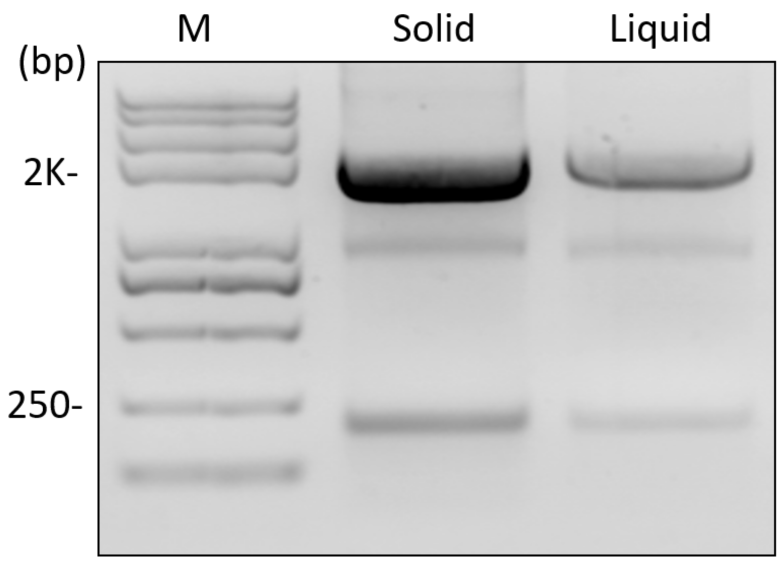


**Figure S9.** Digestion of 3F assembly by NotI (509 oligos pool). M, 2K plus II DNA marker; Solid, cultured on LB plate medium; Liquid, cultured In LB liquid medium.

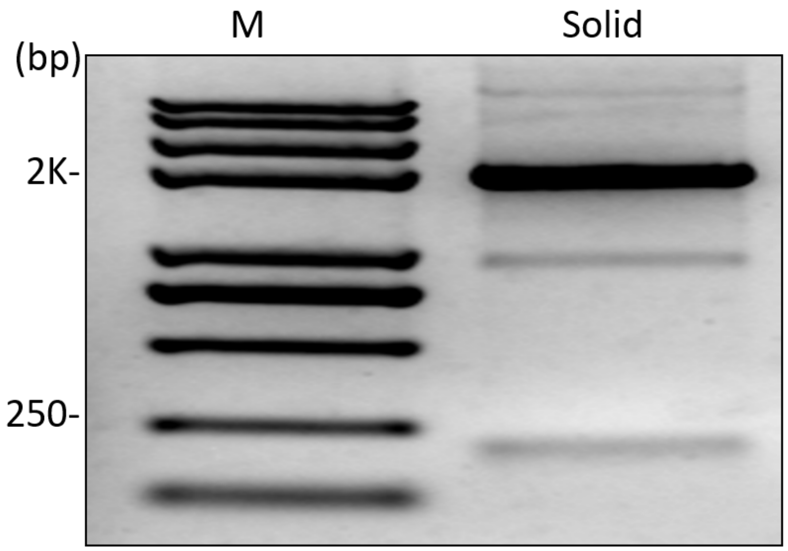


**Figure S10.** Digestion of 5F self-assembly by Not I (509 oligos pool). M, 2K plus II DNA marker; Solid, cultured on LB plate medium.

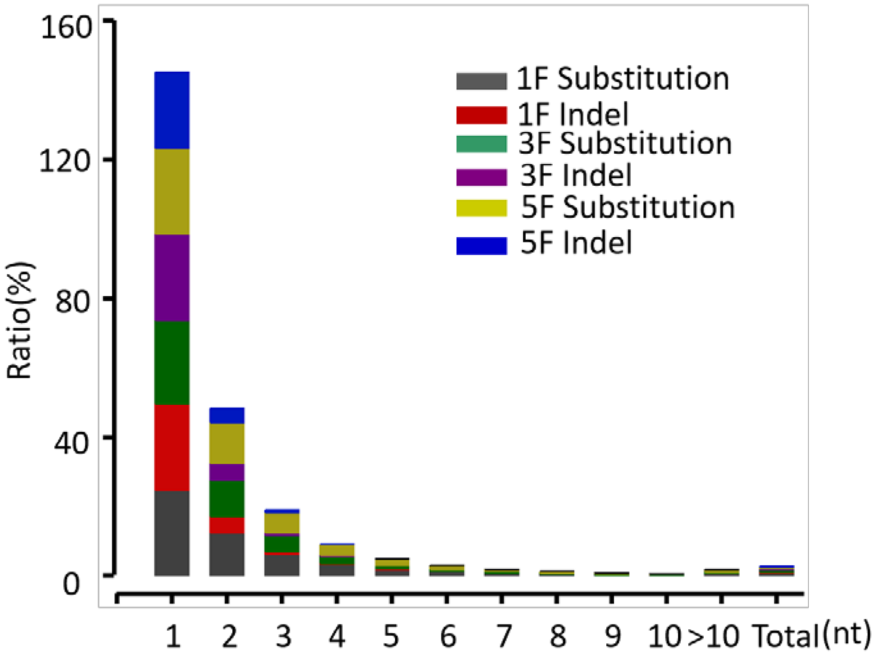


**Figure S11.** Sequenced reads with base error (substitution or indel) were sorted out and the ration of reads with various number of base errors was calculated for assembly of 509 oligos pool.

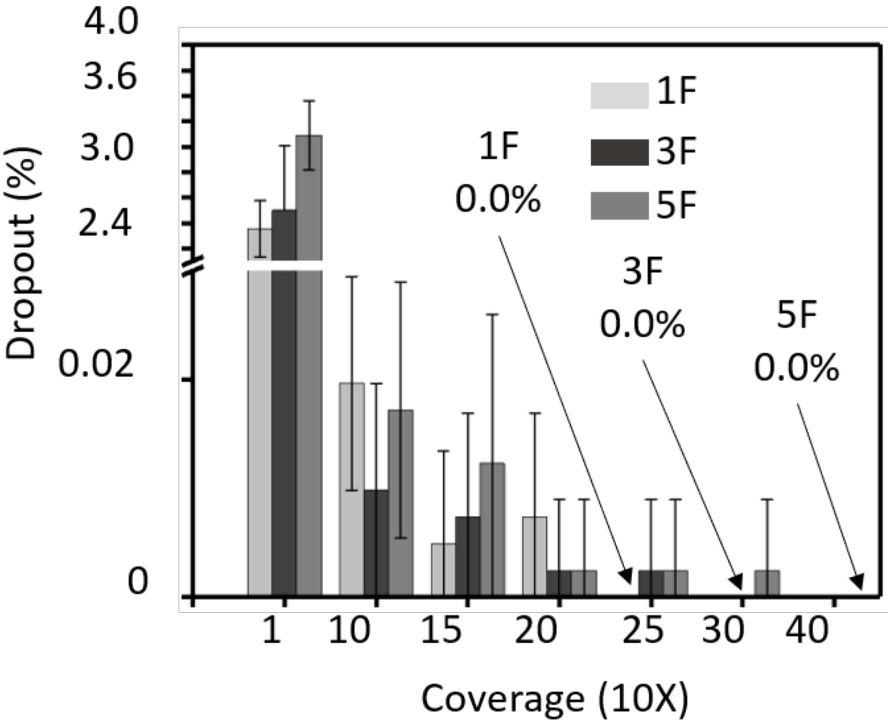


**Figure S12.** The dropout of 1F, 3F and 5F assembly of 509 oligos pool in solid medium culture. When down-sampling the sequencing reads number to 250x of original oligo pool, the dropout of 1F was 0%; When down-sampling the sequencing reads to 300x, the dropout of 3F was 0%; When down-sampling the sequencing reads to 400x, the dropout of 5F was 0%. The position of 0% was indicated by arrow. Error bars represent the mean ± s.d., where n =10.

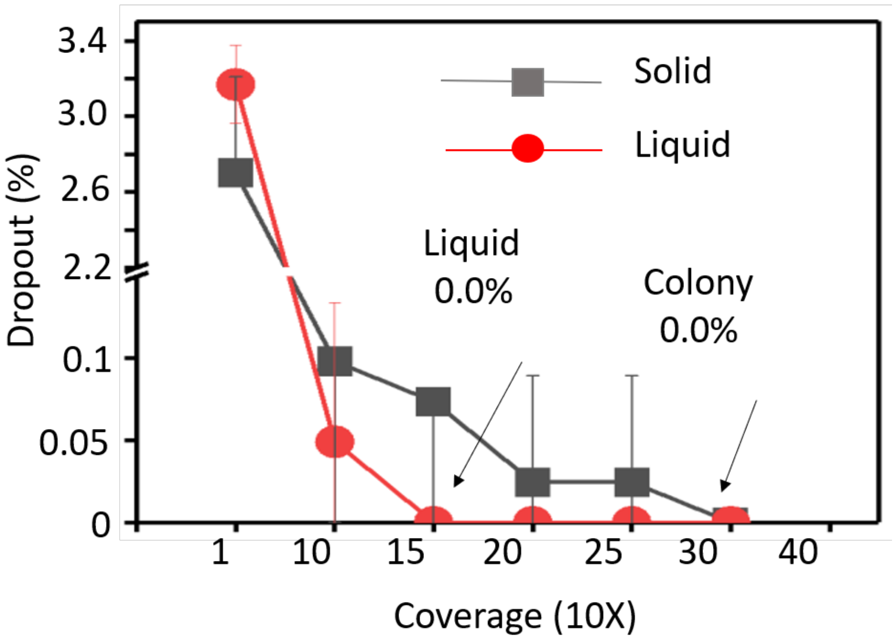


**Figure S13.** The dropout of 3F assembly of 509 oligos pool in solid or liquid culture. When down-sampling the sequencing reads to 150x, the dropout of liquid culture was 0% and down-sampling the sequencing reads to 300x, the dropout of solid culture was 0%. The arrow indicates the position where dropout of each sample is 0 %. Error bars represent the mean ± s.d., where n =10.

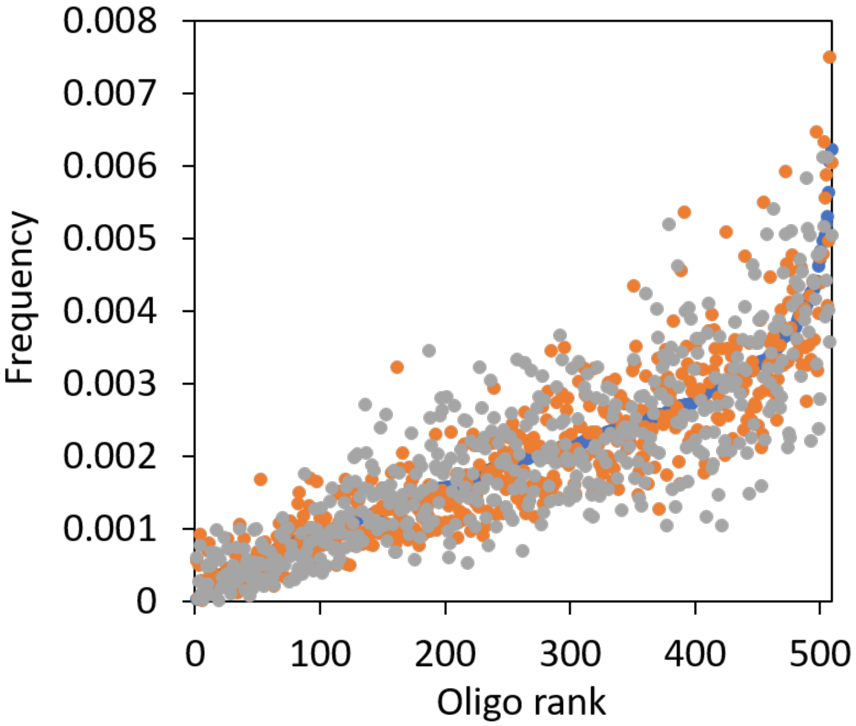


**Figure S14.** Sequenced reads frequency of each 509 oligos reference, for one insert fragment (1F, Blue) assembly, three fragments (3F, orange) assembly and five fragments (5F, gray) assembly.

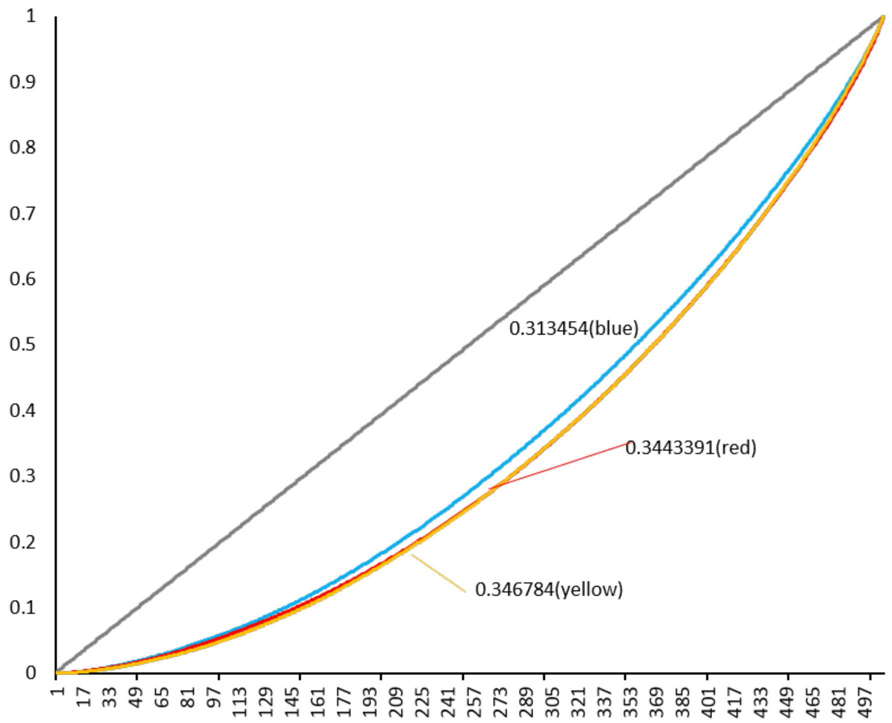


**Figure S15.** Lorenz curve of Gini index for recovered oligos of 1F (blue, with Gini index of 0.313454), 3F (Red with Gini index of 0.3443391) and 5F (Orange with Gini index of 0.346784) assembly of 509 oligos pool.

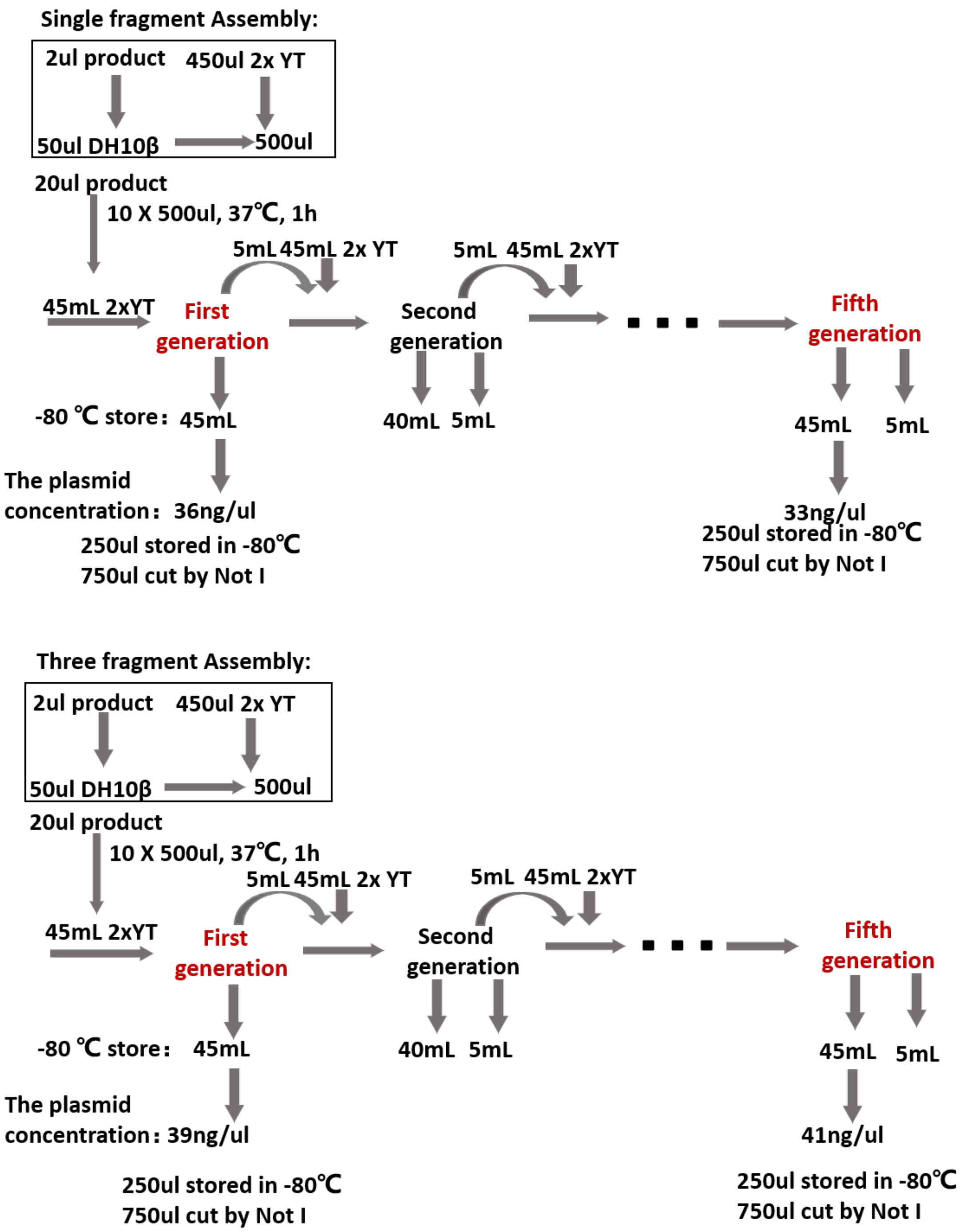


**Figure S16.** The workflow for 11520 oligos pool assembly for mixed culture in liquid medium. The plasmid concentration indicates the extracted plasmid after culture.

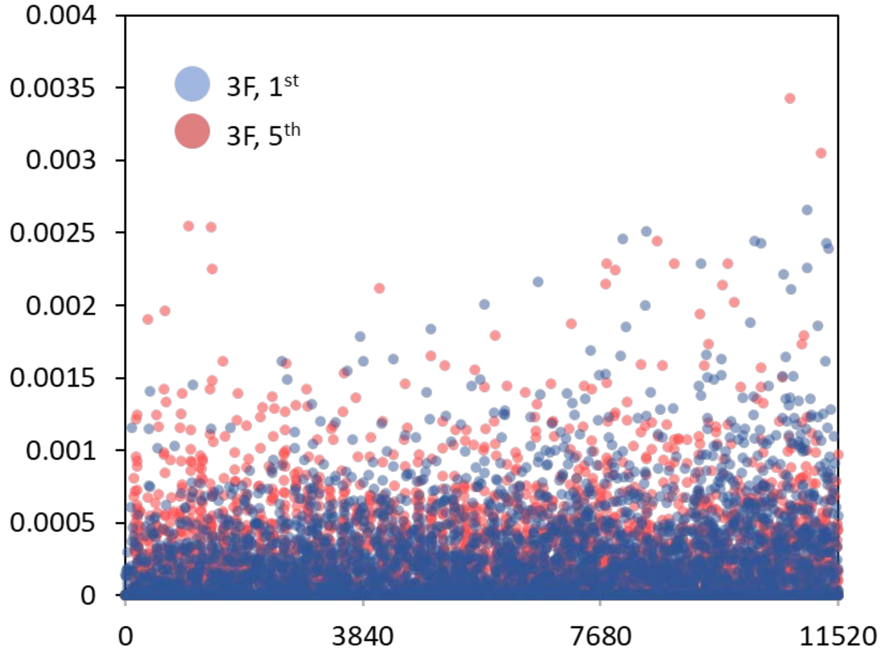


**Figure S17.** The sequenced reads frequency of each 11520 oligos reference of the $1^{st}$ passaging of three insert fragment (3F $1^{st}$, red dot) and the $5^{th}$ passaging (3F $5^{th}$, blue dot).

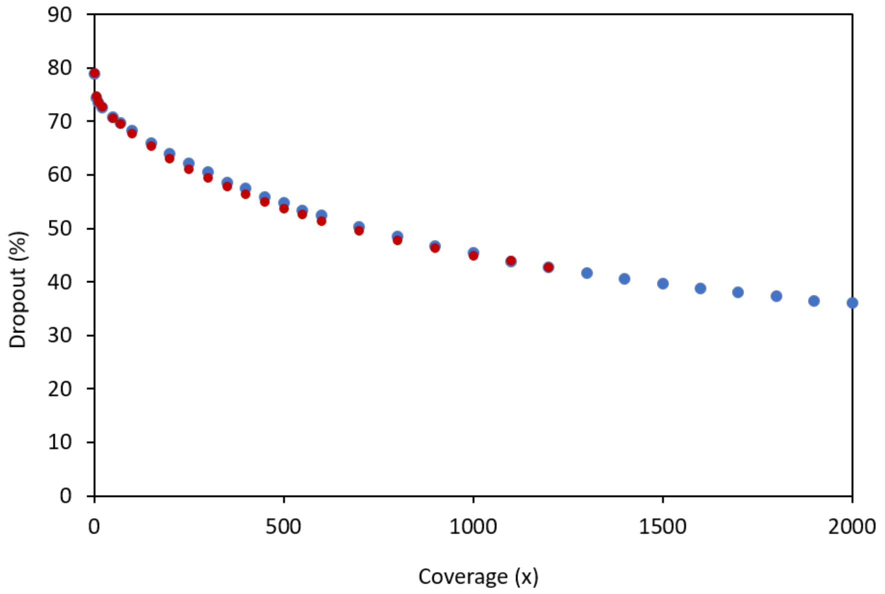


**Figure S18.** Oligo dropout rate was plotted to the corresponding sequencing reads depth. The 1st passaging of three fragment assembly (3F 1st, blue) and the 5th passaging (3F 5th, red).

| Sample | Gini Index |
|---|---|
| Master Pool | 0.29 |
| 1F $1^{st}$ | 0.41 |
| 1F $5^{th}$ | 0.48 |
| 3F $1^{st}$ | 0.87 |
| 3F $5^{th}$ | 0.87 |

**Figure S19.** Gini index of retrieved oligos for each assembled mixed culture of 11520 oligos pool, master pool is the original oligo pool from chip-based synthesis and amplified by PCR.

| Sample | Dropout Rate | Coverage | Perfect Decoding |
|---|---|---|---|
| 1F 1st NotI | 0.90% | 1472x | Yes |
| 1F 5th NotI | 1.40% | 1900x | Yes |
| 1F 1st PCR | 0.40% | 2928x | Yes |
| 1F 5th PCR | 1.40% | 2399x | Yes |
| 3F 1st NotI | 26.50% | 2101x | No |
| 3F 5th NotI | 32.80% | 1325x | No |
| 3F 1st PCR | 25.00% | 2797x | No |
| 3F 5th PCR | 71.70% | 2720x | No |

**Figure S20.** The data statistics for oligo retrieved from 1F or 3F with 11520 oligos pool by direct NotI digest or PCR amplification. The values of dropout rate in the red box are less than 1.56%, corresponding samples can be decoded perfectly. The values in the acid-orange box are over 1.56% and less than 50%, they cannot be decoded. The value in the green box is over 50%, the sample cannot be decoded.

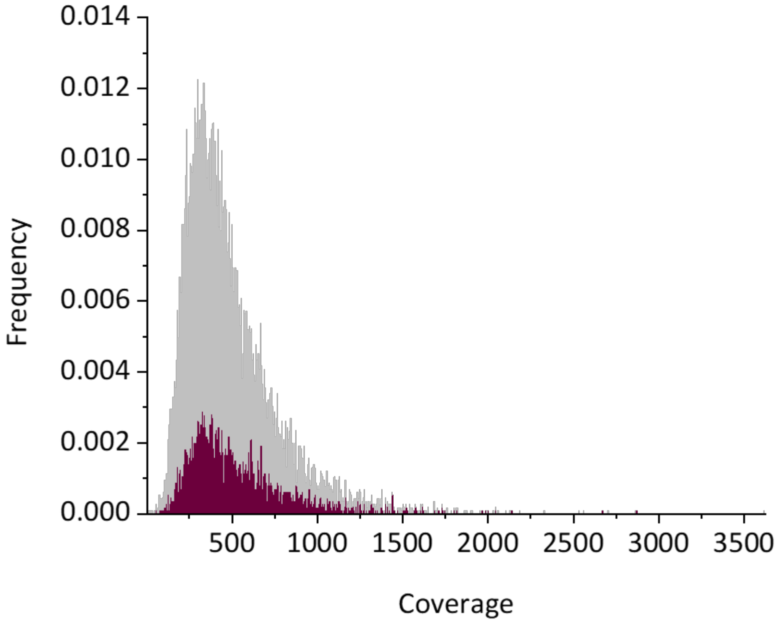


**Figure S21.** The oligo group (red line) which dropout from the 1st passaging of three fragment assembly sample was mapped to the oligos frequency distribution of original master oligo pool (gray line).

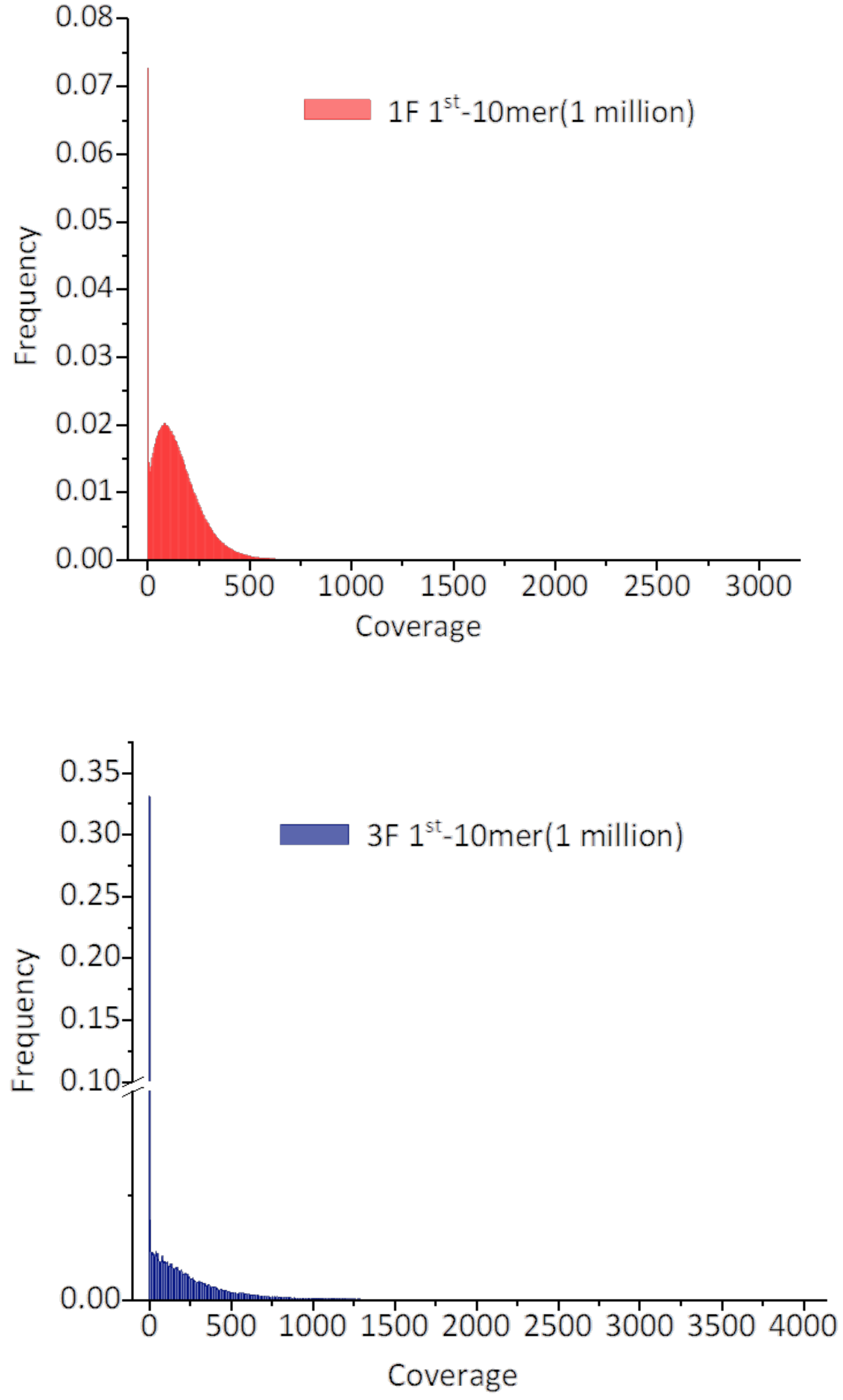


**Figure S22.** 10-mers nucleotide pattern frequency distribution of 1 million valid sequencing reads from sample of 1F 1$^{st}$ (upper) and 3F 1$^{st}$ (Bottom).

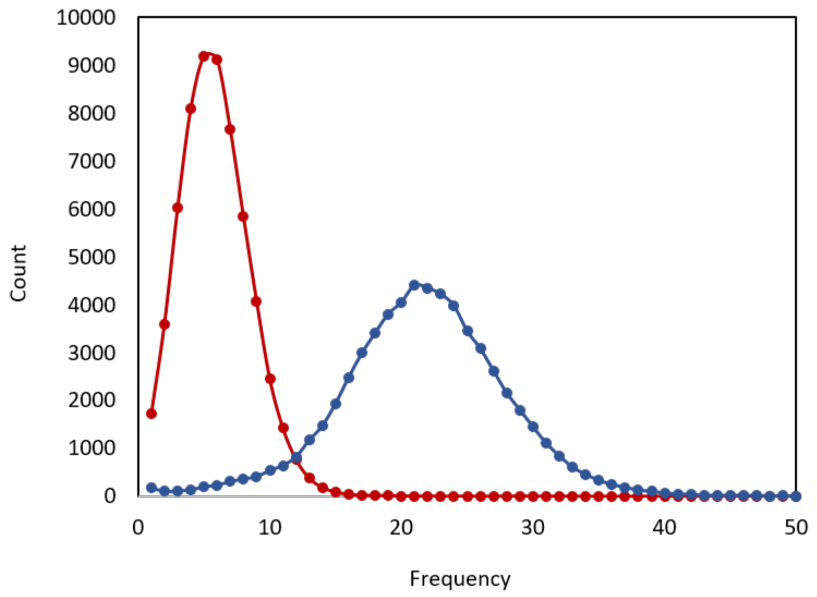


**Figure S23.** 10-mer frequency of three fragment assembly of 11520 oligos pool. Red: enriched oligos, Blue: deprived oligos in comparing with 1F 1st.

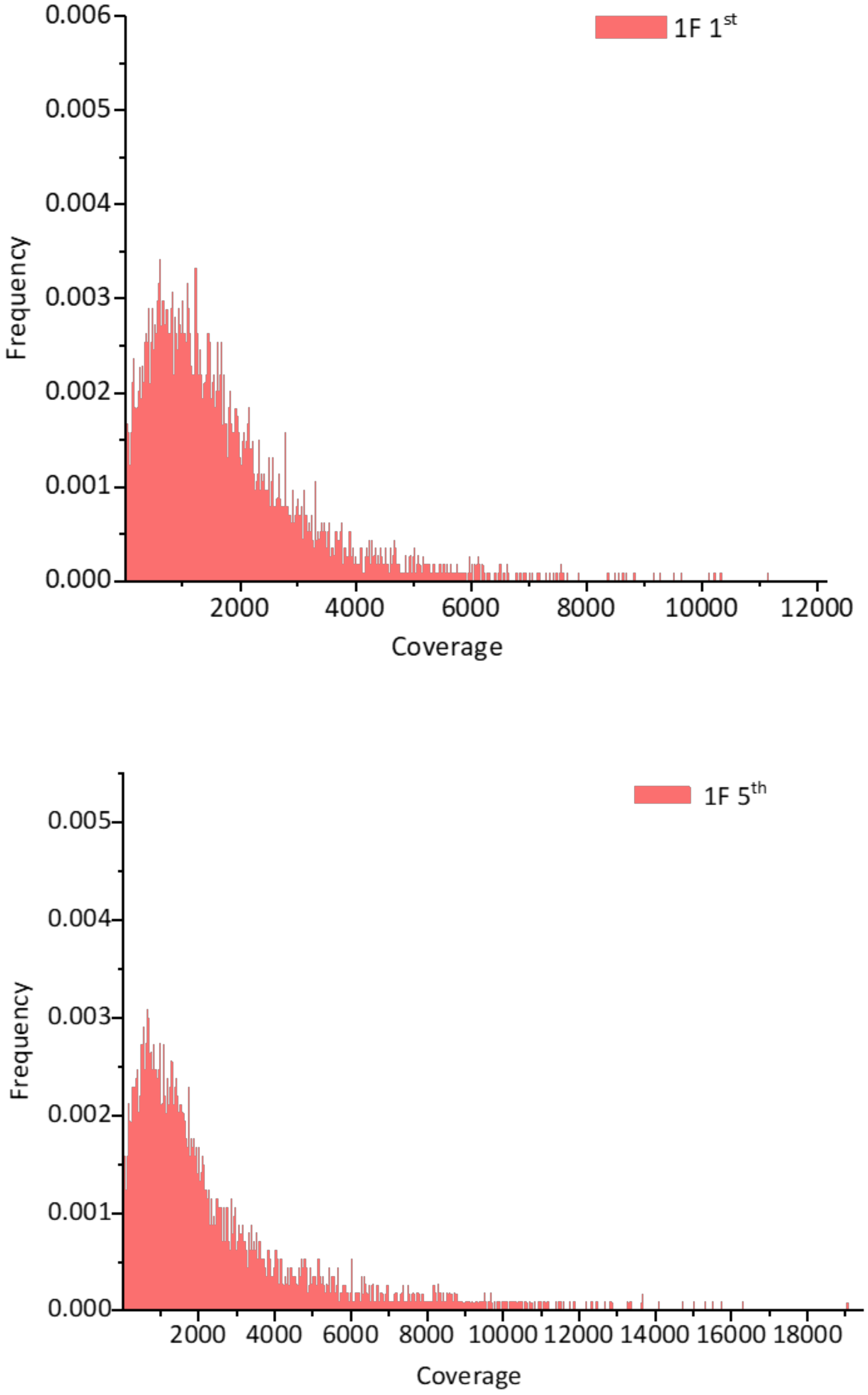


**Figure S24.** Oligo frequency distribution of retrieved oligos from the 1st passaging of one insert fragment assembly (1F 1st, upper) and the 5th passaging (1F 5th, bottom) of 11520 oligos pool.

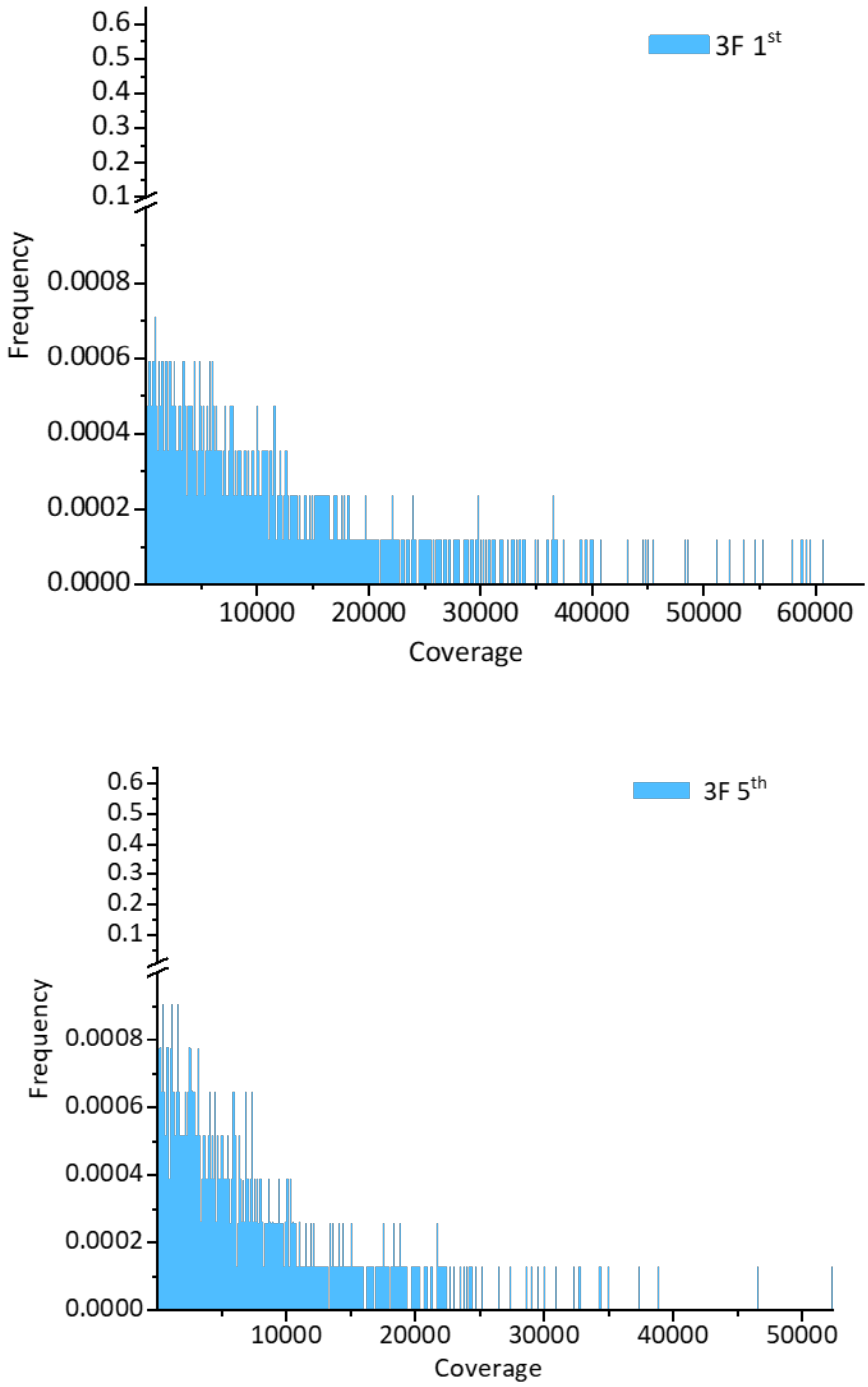


**Figure S25.** Oligo frequency distribution of retrieved oligos from the 1st passaging of three insert fragment assembly (3F 1st, upper) and the 5th passaging (3F 5th, bottom) of 11520 oligos pool.

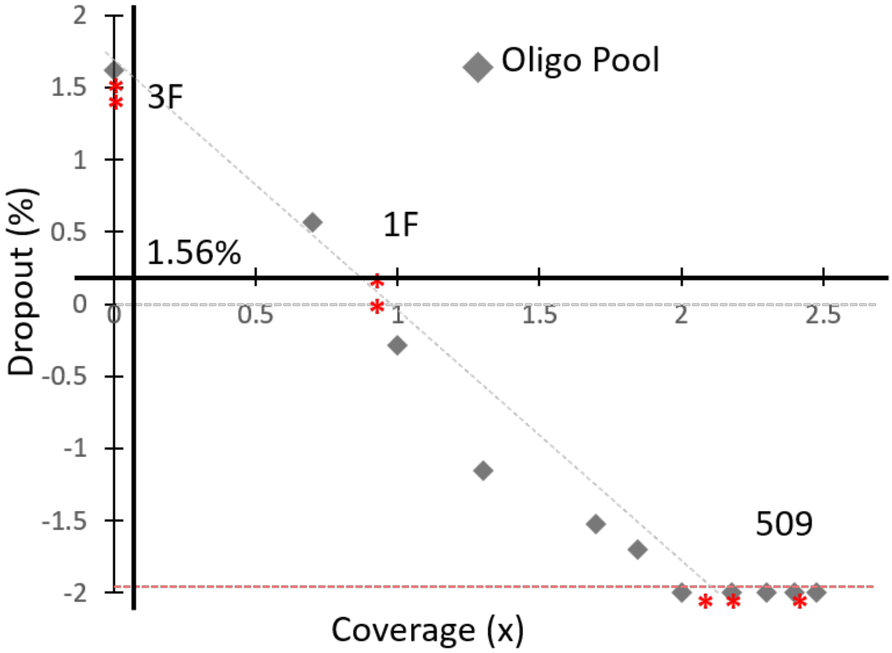


**Figure S26.** Dropout was plotted to corresponding sequencing reads depth. The dropout rate of 11520 master pool was plotted to the corresponding sequencing reads depth (gray diamond) and the dropout rate (red star) of one fragment 11520 oligos assembly sample (1F), three fragments 11520 oligos assembly sample (3F) and all 509 oligos assembly sample respectively were mapped in the dropout rate curve.
